# Local lateral connectivity is sufficient for replicating cortex-like topographical organization in deep neural networks

**DOI:** 10.1101/2024.08.06.606687

**Authors:** Xinyu Qian, Amir Ozhan Dehghani, Asa Borzabadi Farahani, Pouya Bashivan

**Affiliations:** Department of Computer Science, McGill University; Department of Physiology, McGill University; Mila, Université de Montréal

## Abstract

Across the primate cortex, neurons that perform similar functions tend to be spatially grouped together. This biological principle extends to many other species as well, reflecting a common way of organizing sensory processing across diverse forms of life. In the visual cortex, this biological principle manifests itself as a modular organization of neuronal clusters, each tuned to a specific visual property. The tendency toward short connections is widely believed to explain the existence of such an organization in the brains of many animals. However, the neural mechanisms underlying this phenomenon remain unclear. Here, we use artificial deep neural network models to demonstrate that a topographical organization akin to that in the primary, intermediate, and high-level human visual cortex emerges when units in these models are locally laterally connected and their weight parameters are tuned by top-down credit assignment. The emergence of modular organization without explicit topography-inducing learning rules or objective functions challenges their necessity and suggests that local lateral connectivity alone may suffice for the formation of topographic organization across the cortex. Furthermore, the incorporation of lateral connections in deep convolutional networks enhances their robustness to subtle alterations in visual inputs, such as those designed to deceive the model (i.e. adversarial examples), indicating an additional role for these connections in learning robust representations.

## 1 Introduction

Functional organization, characterized by the spatial arrangement of neurons across the cortical sheet according to their similarity in function, stands out as a ubiquitous phenomenon in neuroscience research. This multiscale organization is evident in widely observed topographic maps throughout the brain [HW62, HKPD13, HLB10, OZJ^+^22, GLK^+^18, WKMM78] and has facilitated investigations through lesion experiments, stimulation techniques, and advanced neuroimaging methods such as functional-MRI.

Landmark studies, notably those by Hubel and Wiesel [HW62], examined the micro scale topographical organization of neurons in the visual cortex, with particular emphasis on the structured arrangement of cortical columns based on orientation selectivity and ocular dominance. These observations were later extended to higher stages of the ventral visual pathway with the discovery of neuronal clusters that were selective for faces, scenes, and body parts among others [KMC97, DJSK01, EK98].

The ubiquitous topographical organization of neurons in primate cortices raises two critical questions. First, *why are neurons organized topographically?* A prominent theory addressing this is the Wiring Cost Minimization (WCM) [JJ92], positing that topographical organization is an evolutionary adaptation aimed at minimizing the volume of nerve connections across and within cortical areas. However, it remains unclear whether optimal inter-cortical connectivity has been the sole evolutionary driver of cortical topography. It is possible that cortical topography serves additional functional roles beyond wiring efficiency, yet these roles remain largely unexplored.

Second, *what neural mechanisms enable such multiscale topography across the cortex?* To address the “how” of cortical topography, various computational models have been developed based on the WCM theory. These models employ different strategies including position-aware update rules [Koh82, VdM73, WVDM76], learning objectives that enforce cortex-like position-dependent pairwise response/weight correlations [LMJ^+^20, MLF^+^24, LDB^+^], or penalties on weight connections between distant units [BBP22]. While these models can replicate aspects of cortical topography, such as the formation of category-selective clusters observed in the primate inferotemporal cortex, they exhibit significant limitations.

First, most models only simulate topographical organization in a single brain area (e.g. early or high-level visual cortex) [VdM73, Lin86, BBP22, LMJ^+^20]. Second, models that rely on self-organizing principles often separate the problems of representation learning and topography, assuming sequential or independent mechanisms that are engaged at separate times during development [DM90, DK23]. Third, all topographical models demonstrate diminished performance on ethologically-relevant tasks, such as object categorization [BBP22, DM90, DK23], raising questions about the functional utility of topography. Finally, most models rely on strong assumptions about the underlying biological circuits, such as the need for each neuron to estimate its physical distance to all its input (i.e. presynaptic) neurons [LMJ^+^20, MLF^+^24, BBP22], that are often unmet or remain unspecified.

Substantial evidence suggests that certain aspects of the topographical organization begin to form prior to eye opening. Experiments using animal models such as cats, macaques, and ferrets have shown that ocular dominance columns and long-range correlational structures are present before birth and without visual experience [HH96, LSS78] (see [BS86] for a review). These maps continue to develop postnatally to their mature state [SHW^+^18]. Importantly, sensory deprivation or restrictions could lead to strong degradation in the resulting cortical map [HW97]. Additionally, a large body of literature documents the details of synaptic connectivity across neurons in the primary visual cortex of different animal species [RL82, RL83, CK90, MDCG^+^11, KHP^+^11, KCB^+^13], suggesting that the inherent structure of cortical neural networks plays a critical role in shaping topographical organization.

Two key insights emerge from these studies: 1) Neurons in the primary visual cortex are far more likely to establish lateral connections when they have similar functional selectivity; 2) The likelihood of lateral connections often decays with increasing cortical distance, although there exist patches of distant neurons with strong lateral connections. These insights support two potential interpretations: 1) Neurons with similar selectivity profiles actively establish lateral connections among themselves or 2) Inherent lateral connectivity patterns impose structure on the spatial distribution of neuronal selectivity profiles in the cortex.

In this work, we present Locally-Laterally Connected Neural Networks (LLCNNs), a topographical neural network model that integrates lateral connections into deep convolutional neural networks (Fig. 1). This LLCNN model embodies the hypothesis that local lateral connections alone are sufficient to establish the topographic organization of neurons according to feature selectivity – consistent with the second interpretation above. More importantly, our work reveals a previously unexplored functional role of cortical topography: enhancing the robustness of neural representations. LLCNN consists of a series of convolutional layers with local lateral connections within each layer, representing the reciprocal lateral connections between pyramidal cells, most prominently observed in layers 2 and 3 of the cortex, also known as the superficial patch system [LAB03].

**Figure 1:**
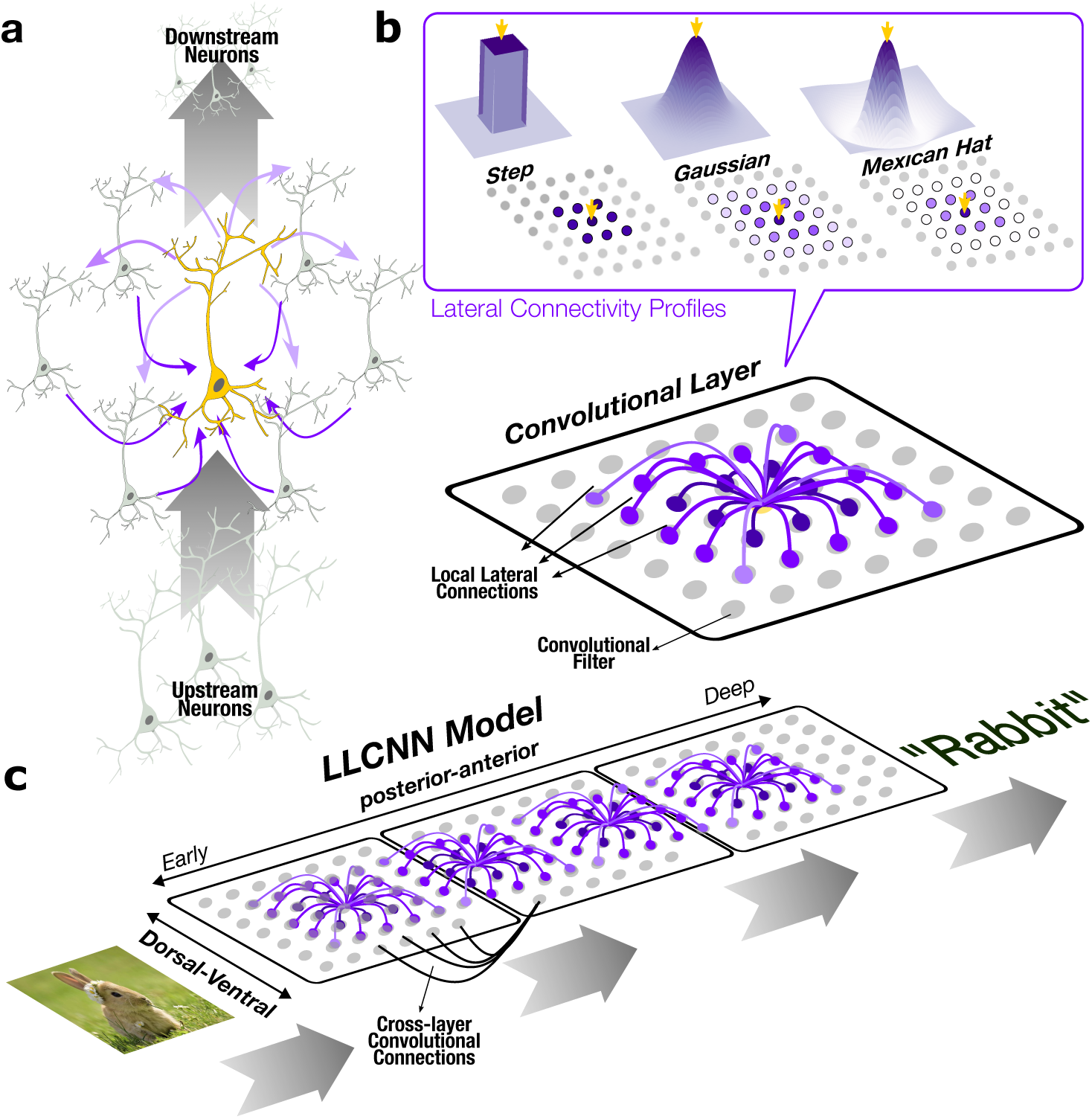
Locally laterally connected neural networks (LLCNNs). a) A schematic of the main circuitry with two types of connections, namely, local lateral connections (in purple) and bottom-up connections from units in an upstream area (in black). b) Convolutional filters are arranged on a two-dimensional “simulated cortical sheet” that defines their spatial configuration. Three variations of distance-dependent connectivity patterns used. Each unit’s output consists of its bottom-up computation (i.e. convolution) and local lateral connectivity whose parameters are fixed and pre-specified as one of three patterns: Step, Gaussian, and Mexican hat. c) Model layers are arranged on a 2D layout with two distinct axes akin to the posterior-anterior and dorsal-ventral axes in the human ventral visual cortex, breaking the permutation symmetry of units in neural networks. Inputs are fed from one end and outputs are generated on the other end.

The locally-laterally connected neural network demonstrates several significant outcomes: 1) It replicates the organization of neurons in the early visual cortex according to the orientation, spatial frequency, and color selectivity; 2) It forms a multi-scale topographical organization in its deep layers according to animacy and object categories, similar to those found in the human inferotemporal cortex; 3) It reproduces the organization in mid-level visual area V4, and predicts unit selectivity in inferotemporal regions with previously uncharted object-selectivity; 4) It improves the trade-off between object recognition performance and cortex-like topography compared to prior models; 5) It shows notable improvement in robustness against adversarial noise, suggesting a previously unknown functional role for cortical lateral connections in learning robust representations.

## 2 Results

### 2.1 Topographical organization of units in early model layers mimics that of V1

Some of the earliest investigations of cortical topography were performed on the primary visual cortex of cats, macaque monkeys and other primates [HW62, BS86]. Broadly, these studies have described the organization of cortex as a periodic map whose periodicity is governed by a number of sensory-dependent features including ocular dominance, eccentricity, visual angle, orientation selectivity, and spatial frequency. Given the abundance of prior experimental work in this area, we first examined whether and to what extent the LLCNN model could replicate the same organization in its early layers that corresponded with the primary visual cortex.

While local lateral connections could in principle be added to any neural network architecture, here we used a commonly used feedforward convolutional neural network model called ResNet18 [HZRS16] that consists of 18 convolutional layers. Units within each layer were arranged on a 2-dimensional sheet simulating a simplified view of the organization of neurons in the cortex (Fig. 1b). Moreover, all layers were also arranged on a 2D layout with two distinct axes akin to the posterior-anterior and dorsal-ventral axes in the human ventral visual cortex (Fig. 1c). This configuration allowed adjacent layers to form lateral connections, similar to the cortical lateral connections that span across distinct visual areas, as previously observed [LH84]. We explored three types of local lateral connection profiles: 1) Step-like function, assigning uniform strength to all connections within a fixed range (LLCNN-S); 2) Gaussian, where connection strengths decay exponentially with distance between units (LLCNN-G); and 3) Mexican-hat, which where connection strengths follows the Mexican-hat pattern with distance between units (LLCNN-MH, Fig. 1b). Besides the addition of local lateral connections connections, all other aspects of LLCNN networks, including training data and procedures, closely matched with conventional methods. All models were trained to perform object categorization using a large dataset of natural images (see Methods).

We evaluated the selectivity of each unit in each model layer (blocks 1 and 2 of ResNet18 architecture) to the stimulus orientation, spatial frequency, and color. For this, we used a stimulus set consisting of gratings with different orientations (0-180 degree), spatial frequencies (1-14Hz), and chromaticity (black/white vs. colored) [MLF^+^24]. We then visualized the selectivity map of units within each layer for each stimulus factor (Fig. 2a). The resulting maps demonstrated smoothly transitioning unit selectivity along the two spatial axes of the simulated cortical sheet for all three stimulus features and for all three variations of LLCNN model (Figs. 2a, S1, S2). We observed that the change in unit selectivity increased to a value of 1 with increasing distance between unit pairs (Fig. 2c), where 1 is the selectivity change expected from random arrangement of units. Likewise, the pairwise response correlation decreased exponentially (Pearson correlation, Fig. 2b). The observed decay in pairwise unit correlation with increasing distance in the early layers of the LLC model suggests that the proximal units exhibit a more congruent response to a set of sine grating stimuli than their distant counterparts, akin to prior observations from the macaque primary visual cortex [NNDC12]. Furthermore, the smooth maps extended across the layers of the LLCNN models, further mimicking the smooth transitioning of these feature selectivity in the brain across cortical areas (e.g. V1 to V2).

**Figure 2:**
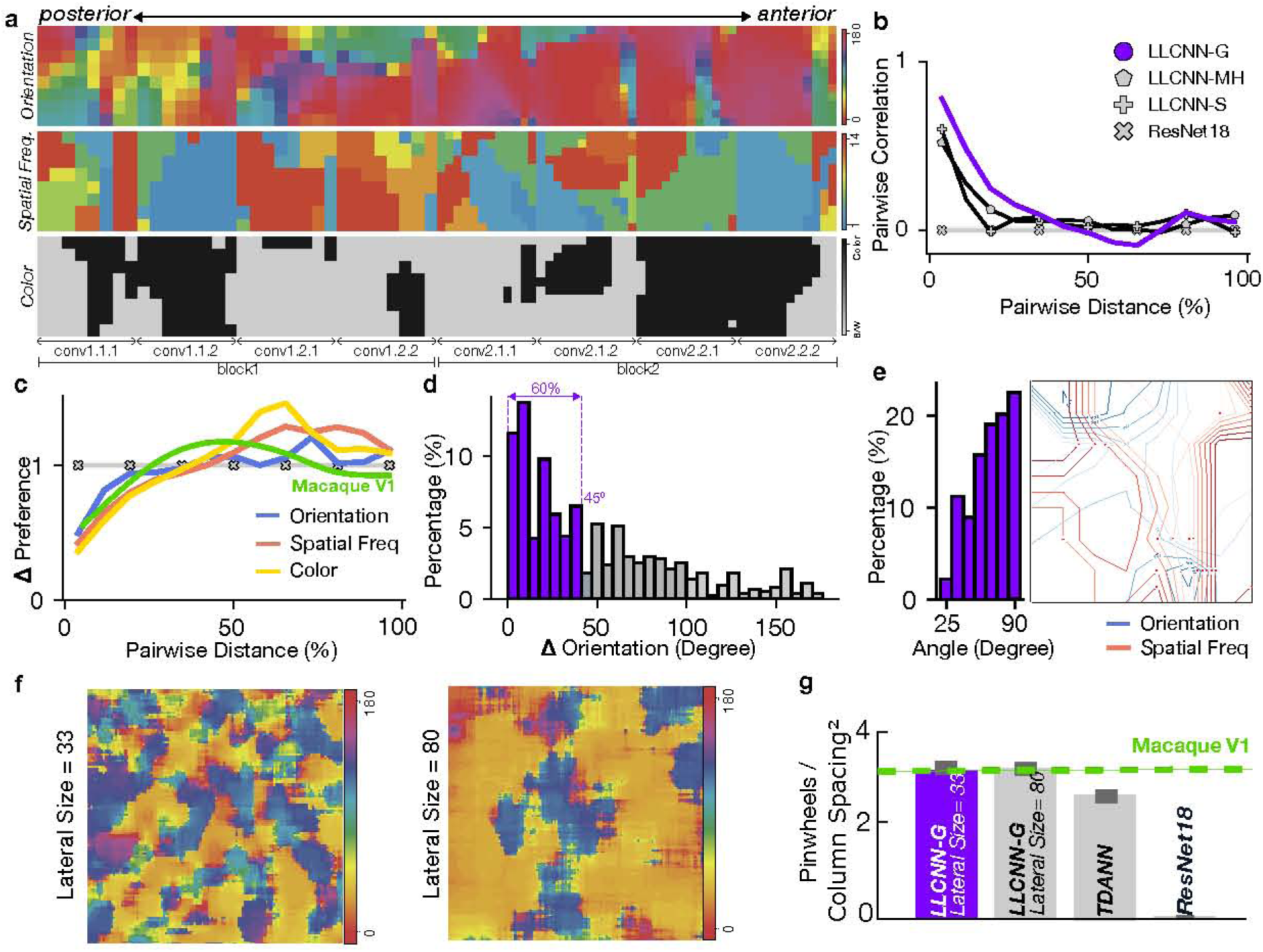
Early LLCNN-G model layers replicate hallmarks of visual processing in the primate primary visual cortex. We first evaluated the topographical similarity of our model with that in the primate V1 by evaluating unit responses to sine grating images of varying orientation, spatial frequency, and color, similar to reference [MLF^+^24]. a) Selectivity maps to orientation, spatial frequency and color smoothly change across the spatial dimensions of the simulated cortical sheet, both within and across layers. b) Pairwise correlation between units decayed exponentially as a function of unit distance on the simulated cortical sheet. Macaque measurements (green curve) were taken from [MLF^+^24] c) Difference in feature selectivity increases and plateaus as a function of distance in an early model layers. d) Distribution of difference in orientation selectivity across immediate neighboring units (i.e. strongly laterally connected units). The proportion of orientation difference ±45^◦^ is ∼60% which aligns with the experimental observation from [MDCG^+^11]. e) A tendency towards orthogonal angles between spatial frequency and orientation gradients similar to prior experimental work [NNDC12]. Orientation (blue) and spatial frequency (red) contours are depicted on the simulated cortical sheet of the first model layer. f) The V1 topography from two variants of locally-connected LLCNN-G with different lateral connection size (Lateral Size = 33 and 75). Typical features of topographical organization in macaque V1 cortex such as pinwheel singularities and linear sections emerge in LLCNN with locally-connected early layer. g) The pinwheel density in locally-connected LLCNN models is comparable with that estimated in macaque V1 (green dashed line; [MLF^+^24]). Larger sigma values yields larger columns and larger relative column spacing, in turn leading to largely stable pinwheel density values (i.e. Pinwheels/Column spacing^2^).

It was also reported that 60-75% of laterally connected neurons in the primary visual cortex of tree shrew [BZSF97], cat [SGLS97], and macaque monkeys [MAHG93] have orientation selectivity that falls within ±45^◦^ of the preferred orientation of the source neuron. We investigated the distribution of difference in orientation selectivity between each unit and its neighboring units and found that 60% of neighboring units have selectivity within 45^◦^ of the center unit (Fig. 2d).

While there are substantial differences in selectivity maps between individuals, these changes still follow particular rules. Notably, using two-photon imaging in macaque’s primary visual cortex, it was shown that the selectivity map gradients (directions on the cortical surface along which the feature tuning changes) of orientation and spatial frequency were markedly skewed towards orthogonality [NNDC12]. We investigated whether the gradients of orientation and spatial frequency selectivity in the LLCNN model followed the same pattern. We computed the gradients of selectivity to each feature and computed the angle between the two gradients at intersection points (Fig. 2e). Our results showed a strong tendency towards crossings at angles close to 90^◦^, echoing the prior findings in the primate primary visual cortex.

Finally, our LLCNN model utilized convolutional layers with shared parameters across different visual spaces. As a result, the model is unable to replicate the retinotopy-dependent organization of the cortex. To explore this aspect further, we developed a modified version of the model where the convolutional layers in the block1 were replaced by locally connected layers. In contrast to convolutional layers, locally connected layers have unique weights for inputs corresponding to different parts of the visual field. After training this model variation, we observed that the orientation preference map of the units in the early model layers displayed a smooth topography, replicating various prominent features of these maps—such as linear sections, singularities, and pinwheels—that are characteristic of the visual cortex [BZSF97] (Fig. 2f). Furthermore, larger lateral connection sizes led to the emergence of larger columns and a reduction in the overall number of columns in the layers. As a result, the pinwheel density in LLCNN-G model variations remained largely stable across and were comparable to that estimated in macaque V1 [MLF^+^24] (Fig. 2g).

### 2.2 Unit arrangement in deep LLCNN layers replicates multi-scale topography in high-level human visual cortex

Neurons demonstrate selectivity for increasingly complex visual patterns as one progresses along the ventral visual pathway. In the high level visual cortex (Ventral Temporal Cortex, VTC), the principle of cortical topography manifests as distinct cortical patches of category-selective neurons, a phenomenon observed across various species, including macaque monkeys and humans. Notable among these are the face-selective fusiform face area (FFA), the body-selective extrastriate body area (EBA) and the scene-selective parahippocampal place area (PPA). Moreover, regions involved in processing specific object classes are organized into partially parallel bands, predominantly running along the posterior-anterior axis of the temporal cortex [BSMT20].

Based on prior literature, we expected the later layers of our model to better correspond to higher visual areas. Consistent with our findings in the early model layers and congruent with prior experimental work [LMJ^+^20], pairwise unit response correlations in these layers also exhibited an exponentially decaying trend as a function of simulated physical distances between units (Fig. 3b). All model variants, regardless of their lateral connectivity function, displayed continuously changing selectivity maps that were significantly smoother than those of the non-topographical model (Fig. 3c). Across different model variations, the model with Gaussian lateral connections (LLCNN-G) demonstrated a more similar decay pattern to that in human.

**Figure 3:**
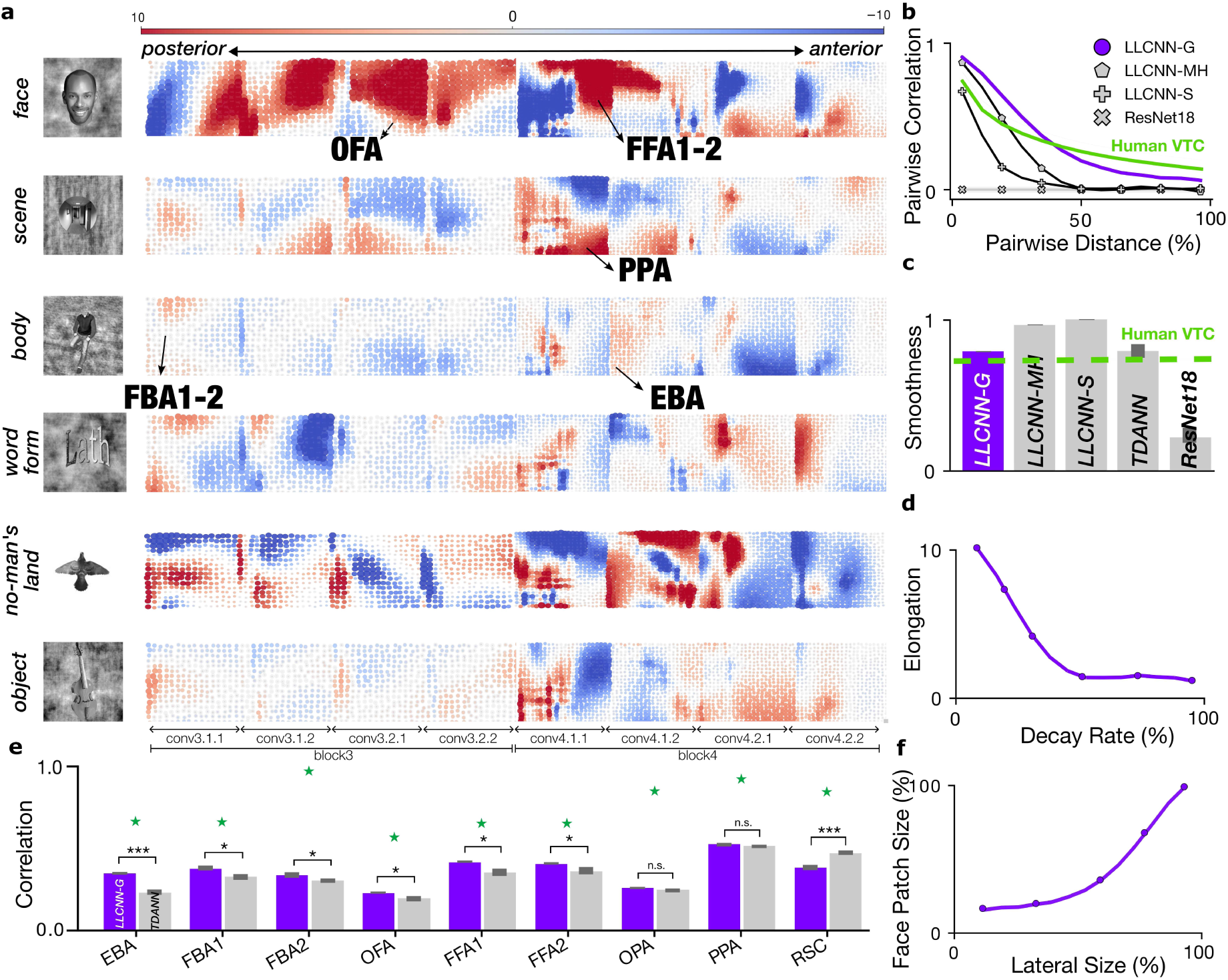
Topograhical arrangement according to object categories in deep layers of LLCNN-G. a) Unit responses were assessed concerning their selectivity to six distinct categories of images, namely face, scene, body, characters, objects, no-man’s land [BSMT20]. The color indicates the strength of selectivity for each category in each unit (red for positive and blue for negative selectivity). Continuous and smooth patches selective for each of the six categories emerged in the deeper layers of the model (block 3-4) that were significantly elongated along the posterior-anterior axis of the model, similar to typical elongation of category selective patches along the posterior-anterior axis of ventral visual cortex [BSMT20] b) Pairwise unit correlations decayed exponentially as a function of distance. Human VTC curve was calculated from the NSD dataset [ASYW^+^22]. c) LLCNN-G yielded a smoother topography compared to other variants and comparable to TDANN. d) The patch elongation decreases as a function of how fast the lateral connection range was decayed during training. e) Activity in category-selective LLCNN-G patches closely resembled those in previously studied category-selective cortical patches. Comparisons were made using the NSD dataset [ASYW^+^22] and error bars indicate the standard deviation across subjects. Green stars indicate the internal consistency of the brain activity signal in each region. f) Patch sizes were positively modulated by the size of lateral connections.

We then examined whether units in deeper layers of our neural network similarly clustered according to their category-selectivity as in the human brain [KMC97, EK98, DJSK01]. We quantified selectivity of each model unit to six distinct image categories–face, scene, body, characters, objects, no-man’s land [ASYW^+^22, BSMT20]–and visualized them on the model’s simulated cortical sheet (Fig. 3a). We identified distinct unit clusters for each category in the deeper layers of the network (layer 3 and 4 of ResNet18 architecture). Pairwise response correlations decayed exponentially as a function of distances between units in all model variations (Fig. 3b) and the unit selectivity changed smoothly along the spatial dimensions of the simulated cortical sheet (Fig. 3c), comparable to previous model of cortical topography TDANN [MLF^+^24]).

In addition, most category-selective clusters were significantly elongated along the posterior-anterior axis of the simulated cortical sheet, forming semi-continuous bands that expanded over multiple layers of the network (Fig. 3a). This feature closely mirrors the relative elongation of similar patches in the human and monkey visual cortex [NLD^+^11, BSMT20]. Additionally, the extent of this elongation along the posterior-anterior axis varied with the decay rate of the lateral connectivity window size during training, with faster decay rates leading to less elongation (Fig. 3d). Training the LLCNN model with a fixed lateral connection window size resulted in blob-like category-selective patches with milder elongation effect along the anterior-posterior axis (Fig. S7). Similarly, the size of category-selective patches co-varied with the size of the lateral connectivity window (Fig. 3f). Similar to the human cortex, face- and scene-selective were strongly anti-correlated and units with selectivity to each category formed notably non-overlapping pathways positioned on the opposite sides of the dorsal-medial axis of the simulated cortical sheet (Fig. 3a). The highest model patch similarity to OFA appeared in layer 3, preceding the FFA, which emerged in layer 4 (similar patterns were observed for FBA-1 and EBA regions). For other object categories, while multiple selective patches appeared across the model layers, they did not form clear bands like those for face and scene processing. Interestingly, apart from face and scenes, selectivity to NML category was among the highest and most widespread among the six categories. Finally, the activity in LLCNN-G category-selective patches were highly correlated with those measured from different category selective human cortex, improving upon TDANN model in several cortical areas including OFA, FFA, and EBA (Fig. 3e). We observed that training the LLCNN models using an unsupervised learning objective [ZJM^+^21] resulted in qualitatively similar results (Fig. S6). However, we didn’t perform quantitative comparisons between supervised and unsupervised trained models.

Having investigated the topographical organization of the model at mesoscale, we next explored whether it also replicated the macroscale organization observed in the human visual cortex. We analyzed unit arrangements according to their selectivity for object animacy and real-world size, extensively investigated in human studies [KO12, KC13]. Consistent with human observations, model units clustered according to preferences for animate versus inanimate objects and for small versus large real-world objects. Within each map (animacy and size), units with similar selectivity formed parallel streams along the posterior-anterior axis of the model (Fig. 4a). This finding differed significantly from those previously reported in self-organized topographical neural network models [DK23] but closely resembled the extended arrangement of these pathways along the posterior-anterior axis in the human ventral visual cortex [KC13].

**Figure 4:**
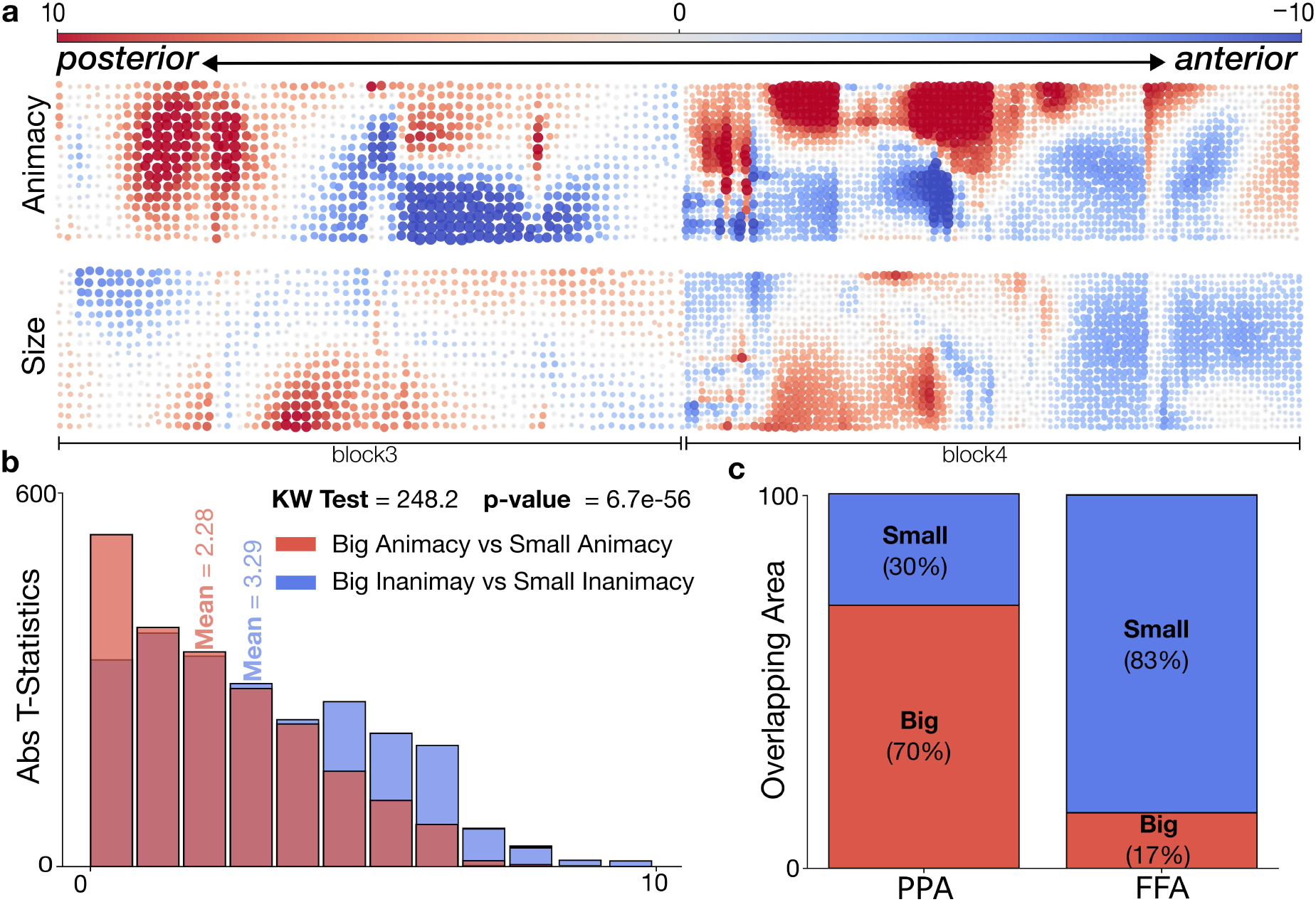
Macro-scale organization based on animacy and object size emerges across LLCNN-G layers. a) Unit selectivity to animate vs. inanimate objects, as well as, large vs. small objects emerge along continuous bands. The model displayed two parallel streams that encoded animacy and size of objects, similar to prior observations from human visual cortex. Plot shows the 8 convolutional layers in Blocks3 and 4 of the model. b) The t-statistics for big animacy vs. small animacy and big inanimacy vs. small inanimacy show a statistically significant difference between the means of these two sets of t-values. c) The overlapping area between size and PPA/FFA reveals that the PPA overlaps mostly with patches that are selective to big size (70%), while the FFA mostly overlaps with patches selective to small size (83%). [KC13]

Moreover, it has been shown that the similarity between small and big animals is significantly greater than that between small and big objects [KC13]. We investigated whether these differences in selectivity also existed in the LLCNN model units. For this, we calculated each unit preference for big or small objects using the T-statistic and contrasted the distribution of selectivity for size as a function of unit’s preference for animate or inanimate objects (Fig. 4b). The result showed that LLCNN-G units that are selective to animals were significantly less sensitive to size, compared to those that are selective for inanimate objects. In addition, we found that the 70% of units in the scene-selective PPA patch of the LLCNN-G model were selective for big objects, while 83% of units in model’s FFA were selective for small objects (Fig. 4c), further replicating prior observations of similar overlaps in the human cortex [KC13].

### 2.3 Topographical organization in uncharted and non-categorical cortical landscape

We further investigated whether the representations in intermediate layers of the LLCNN model also correspond to the intermediate visual areas in the primate brain. Neurons in macaque monkeys’ area V4 were previously shown to form clusters according to their selectivity to corner or curvature stimuli [JALT21]. We examined the selectivity of units in LLCNN model to corners and curvatures using the same stimulus set used in prior work. We found that the units in block 3 of the LLCNN model were similarly clustered according to their preference for corner versus curvature stimuli (Fig. 5a). Furthermore, similar to our previous observations in early and deep layers of the model, the pairwise unit correlations exponentially decayed as a function of distance between units (Fig. 5b). The selectivity for curve or corner patterns also followed a similar pattern to experimental data from macaques (Fig. 5c; [JALT21]). Visual examination of the best stimulus maps for stimuli from alternative stimulus sets [WRF^+^23, PC02] further confirmed the existence of robust unit clusters and smooth transition in selectivity across the model units (Fig. S8).

**Figure 5:**
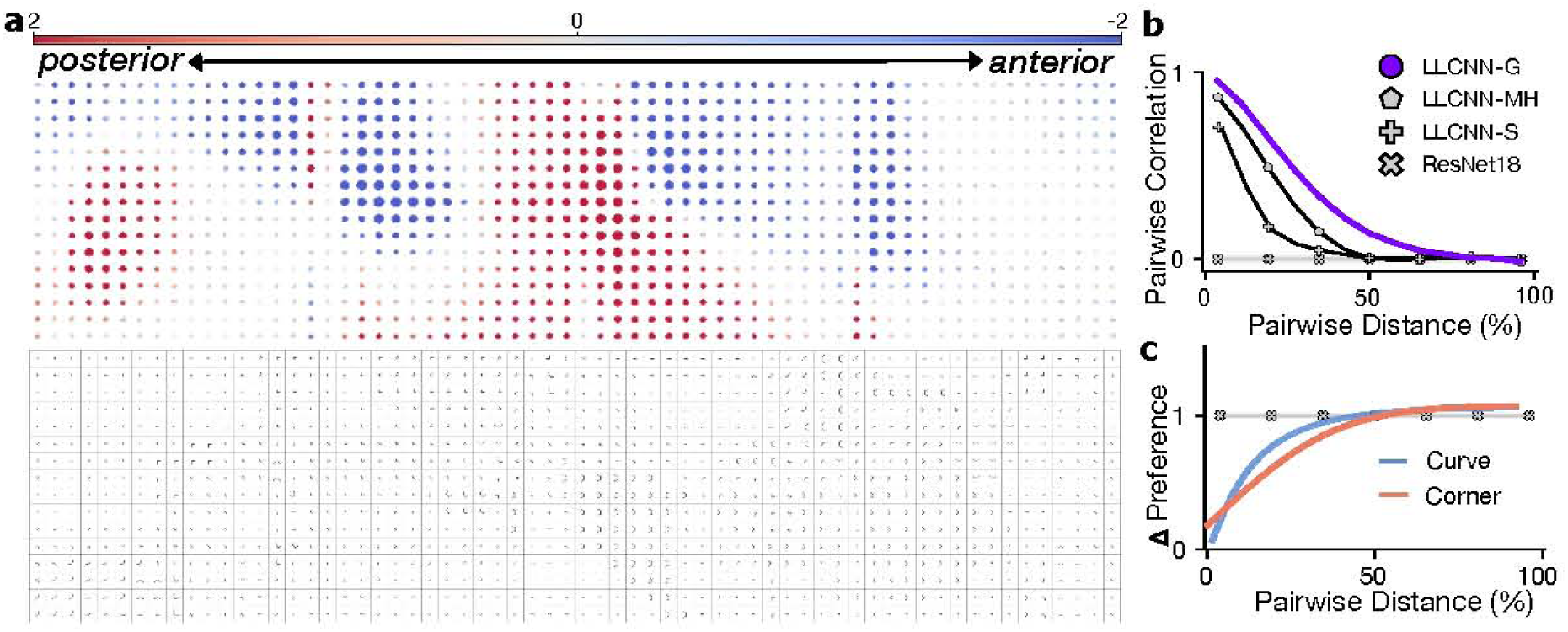
Topographical organization of units in intermediate layers of the LLCNN-G model according to selectivity to corners and curves. a) We investigated the V4-like topographical organization in block 3 of the LLCNN-G model by examining the unit selectivity for curves and corners. (top) t-statistics are shown for individual units in the 4 layers in block 3 of the model. (bottom) best stimulus from stimulus set used in [JALT21] eliciting the highest activity in each unit is shown. b) The pairwise correlation of unit activations responding to shape dataset decayed exponentially as a function of pairwise distance. c) The units’ selectivity to corners and curves increased exponentially as a function of pairwise physical distance.

We next asked whether the topographical organization of our model also aligns with that in the human cortex outside of the category-selective regions. To perform this analysis, we first identified the center point of FFA-1 and PPA regions that are most consistent across individuals. We next designated the intermediate point between the FFA-1 and PPA center points in each subject as the *intermediate* patch (Fig. 6a). Our hypothesis was that if the model’s organization is a good match to that in the human visual cortex, the activity in this intermediate patch should be well correlated with the activity in units positioned at the intermediate point between the model’s respective patches. We calculated the Pearson correlation between all LLCNN-G units and each patch in the brain. The resulting correlation maps showed that indeed the units that were most highly correlated with the intermediate brain patch were located at spatial positions in between the units highest correlated with FFA-1 and PPA (Fig. 6b). All LLCNN model variations could predict the brain activity at the intermediate patch relatively well while the Gaussian variation was significantly better than the other two variations (Fig. 6c). This result suggested that the unit arrangement in these models is not only aligned with the brain in terms of replicating the category-selective patches but also generalizes to uncharted cortical landscape in between.

**Figure 6:**
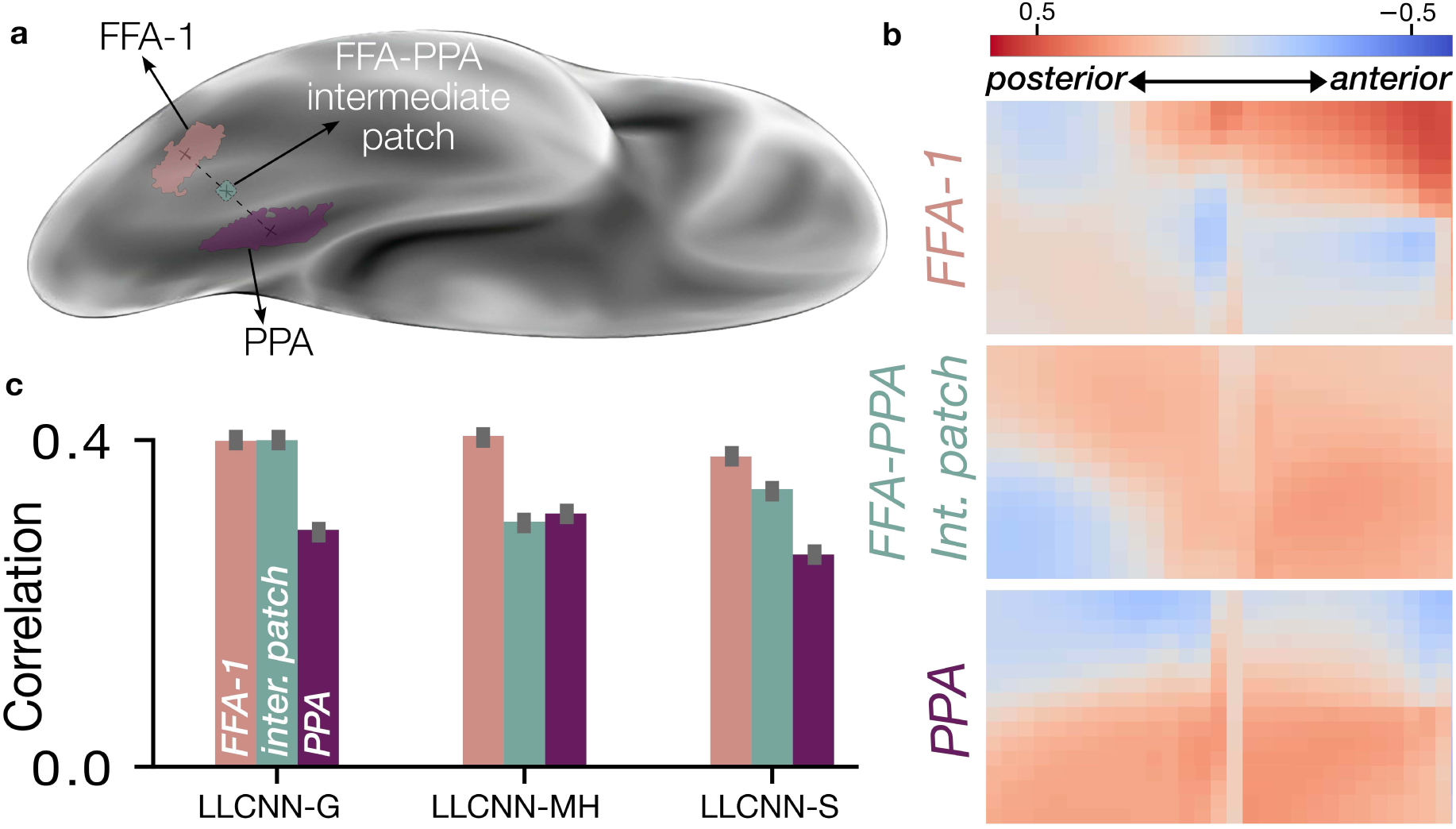
Predicting activity in uncharted parts of the human VTC cortex. a) Identifying an intermediate cortical patch between FFA-1 and PPA in the right hemisphere from NSD dataset (subject 1 is shown). b) Correlation map for the LLCNN-G model units in block 4 and each patch in the brain. Units with the highest correlation to the intermediate patch fall (spatially) predominantly in between the units with maximum correlation with FFA-1 and PPA. c) Correlation between intermediate LLCNN-G unit and the intermediate cortical patch for different model variations. Error bars indicate the standard deviation across subjects.

### 2.4 Impact of local lateral connections on learned behavior and internal representations

#### Performance on visual categorization

How does local lateral connectivity affect behavioral performance? Prior work on topographical neural network models had reported a conflict between topography-inducing objective functions and task learning (e.g. visual categorization) [MLF^+^24]. In this perspective, the parameter updates necessary for learning better representations in practice clash with those necessary for organizing units in a topographical organization, leading the training dynamics to reach an equilibrium that reflects the trade-off between the two objectives. The magnitude of this impact on visual categorization is often large and undesired (∼25.7% reduction on Imagenet accuracy in TDANN [MLF^+^24]). We evaluated the categorization performance of different LLCNN variants and compared them with non-topograhpical ResNet18 and a recent topographical neural network model [MLF^+^24]. Although LLCNN-G model displayed a significant drop (16.5%) in object recognition performance compared to its non-topographical counterpart, its accuracy was still substantially higher than the state-of-the-art topographical model [MLF^+^24] (TDANN=43.9%, LLCNN-G=53%, RN18=69.57%; Fig. 7c)^1^.

**Figure 7:**
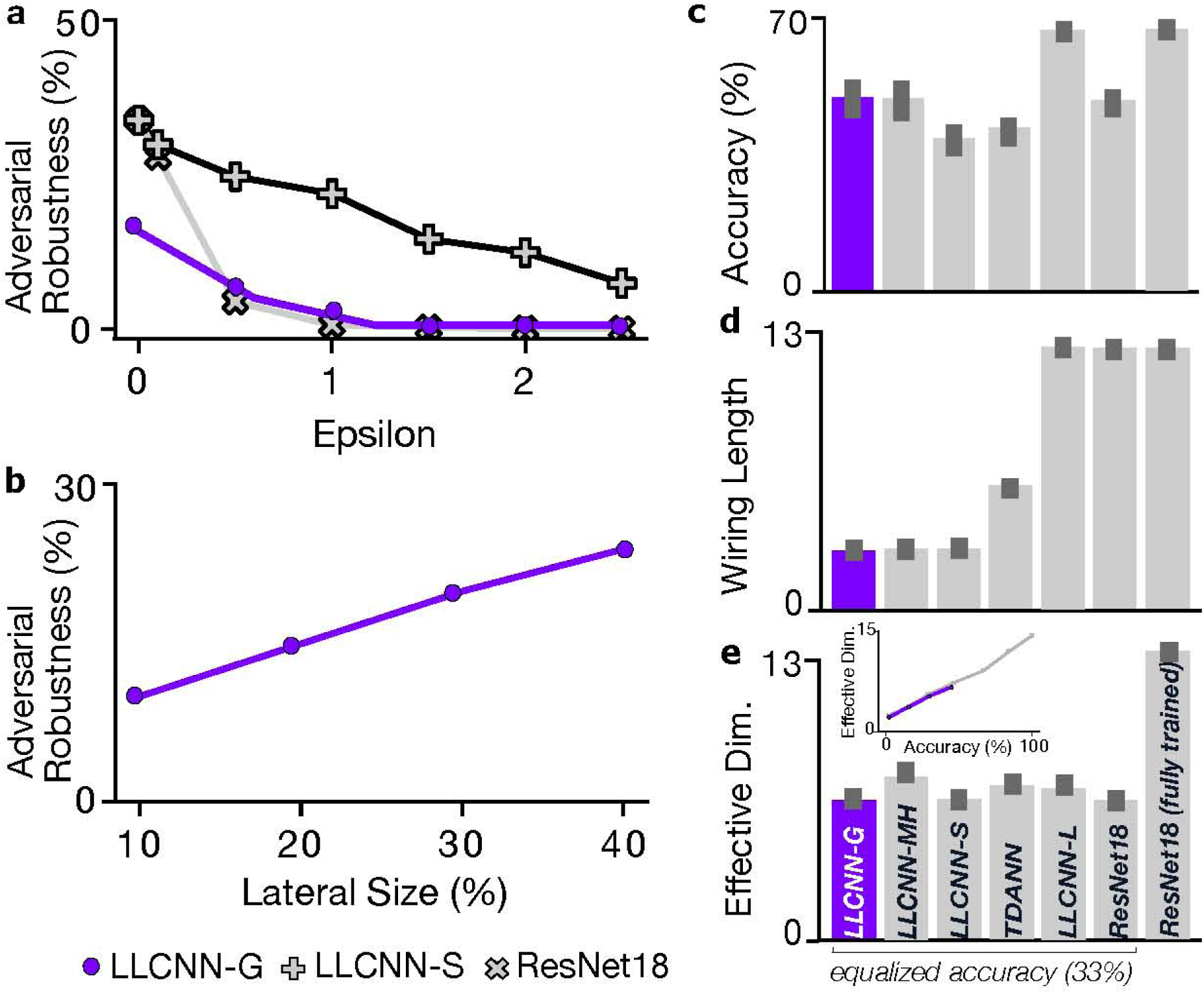
Behavioral performance and Robustness. We compared the performance of the LL-CNN with that of baseline models. a) LLCNNs show stronger resilience to adversarial perturbations (AutoAttack-L2) compared to the non-topographical model ResNet18. Epsilon denotes the strength of the allowed image perturbation measured with *L*_2_ norm; b) Larger lateral size yields larger robustness to adversarial perturbations. Adversarial robustness was evaluated using AutoAttack *ɛ*_L2_ = 0.5; c) LLCNN-G and LLCNN-MH exhibit better trade-off between visual categorization accuracy and topography compared to other baselines. d) Training LLCNNs with gradient descent leads to minimization of the total wiring length in the network. e) The effective dimensionality does not significantly differ between topographical and non-topographical models with equalized accuracy (33%). Inset shows the relation between effective dimensionality and object recognition accuracy on ImageNet dataset for intermediate checkpoints of ResNet18 (gray) and LLCNN-G (purple) during training. Effective dimensionality linearly increases with improved accuracy for both models.

#### Connection lengths

According to the WCM theory, the primary role of the modular organization in cortex is to minimize the volume of interconnecting nerves. We next investigated whether inclusion of local lateral connections results in the reduction of connection lengths between layers and to what degree. We used a previously used metric[MLF^+^24] to estimate the global connection weights in the network from the connection weights. Using this metric, we found that the wiring length of LLCNNs is significantly lower than the non-topographical ResNet18 model as well as another recent topographic model that was explicitly optimized for this objective [MLF^+^24]. In fact, the wiring length in LLCNN-G was 16.7% of the non-topographical one and about half of that in TDANN (LLCNNG=2, TDANN=6, ResNet18=12; Fig. 7d). This suggests that top-down learning in neural networks (e.g. via backpropagation) with local lateral connectivity on ethologically-relevant tasks such as object recognition minimizes the wiring cost as a byproduct. It surprisingly does so even more effectively than alternative topographical models that induce topography by enforcing brain-like distance-dependent correlation structure between model units [MLF^+^24].

#### Robust representations

We posited that the learned unit clusters in LLCNNs may be implementing a form of ensemble model where each feature value is computed by finding the average estimate of multiple units with similar selectivity. Machine learning ensemble approaches such as Bagging and Boosting were previously shown to be robust to input perturbations [Fre95, Bre96, Bre01, HTFF09]. We tested the LLCNN models to verify if such robustness to input perturbations is also present in the models by subjecting each model to input images that were adversarially perturbed using a strong adversarial attack called AutoAttack [CH20]. The neural network with lateral connections displayed strong resilience to pixel perturbations (Fig. 7a) compared to the non-topographical model. While the performance of both models eventually reached zero with larger perturbations (i.e. epsilon denoting the magnitude of the perturbation measured with *L*_2_ norm), the LLCNN model with step-like local lateral connectivity showed consistently higher robustness compared to the topographical one. Furthermore, we observed that robustness increases with larger lateral connection range in the model (Fig. 7b; AutoAttack *ɛ*_L2_ = 0.5). This suggested a possible auxiliary role for lateral connections in the cortex in learning robust representations.

#### Effective dimensionality

Lower effective dimensionality has previously been reported in topographic models of visual cortex and was proposed as an inherent characteristic of these models that leads to better alignment with the neural responses in the brain [MLF^+^24]. We found that while the effective dimensionality of all topographical models were substantially lower than the original ResNet18 architecture, this difference becomes insignificant for most models, when the categorization performance of these models is matched (Fig. 7e). Without controlling the accuracy of ResNet-18, there is a positive correlation between complexity and accuracy that may lead to higher dimensionality in ResNet-18 with maximum accuracy. This result suggests that the significantly lower-dimensional representations in topographical models is a byproduct of their lower categorization accuracy rather than an inherent feature.

To gain further insight into the underlying representation in the LLCNN-G model, we examined the unit activity profiles [DK17] in a category-independent manner. For this, we computed the principal components of the transposed activation matrix (i.e. in which units and stimuli are arranged along the rows and columns respectively) in response to the NSD stimulus set. In this representation of activations, each sample represents the specific unit’s activity profile to the complete set of stimulus set, making the PCA space a proxy for the function space learned by units in LLCNNs.

We found that units in non-topographical ResNet18 span a sphere-like function manifold showcasing diverse and uniform response profiles across the units. In contrast, the function manifold in the LLCNN-G model is much lower dimensional, showing distinct topological properties (Fig. 8a). By coloring the units based on their physical positions in LLCNN-G model, we observed that the manifold preserves the pairwise relative distances in the model, indicating that units in close spatial proximity tend to produce more similar responses. Similar structure existed in block3 of the model as well (Fig. S9). Similarly, repeating this analysis on brain activity using the NSD dataset, we observed a similar smooth and low dimensional manifold in the human VTC cortex (Fig. 8b). Moreover, most human VTC cortex PCs were significantly more aligned with PCs from the LLCNN-G model compared to those from non-topographical ResNet18 (Fig. 8c). Nevertheless, we could not identify any significant semantic meaning associated with these directions.

**Figure 8:**
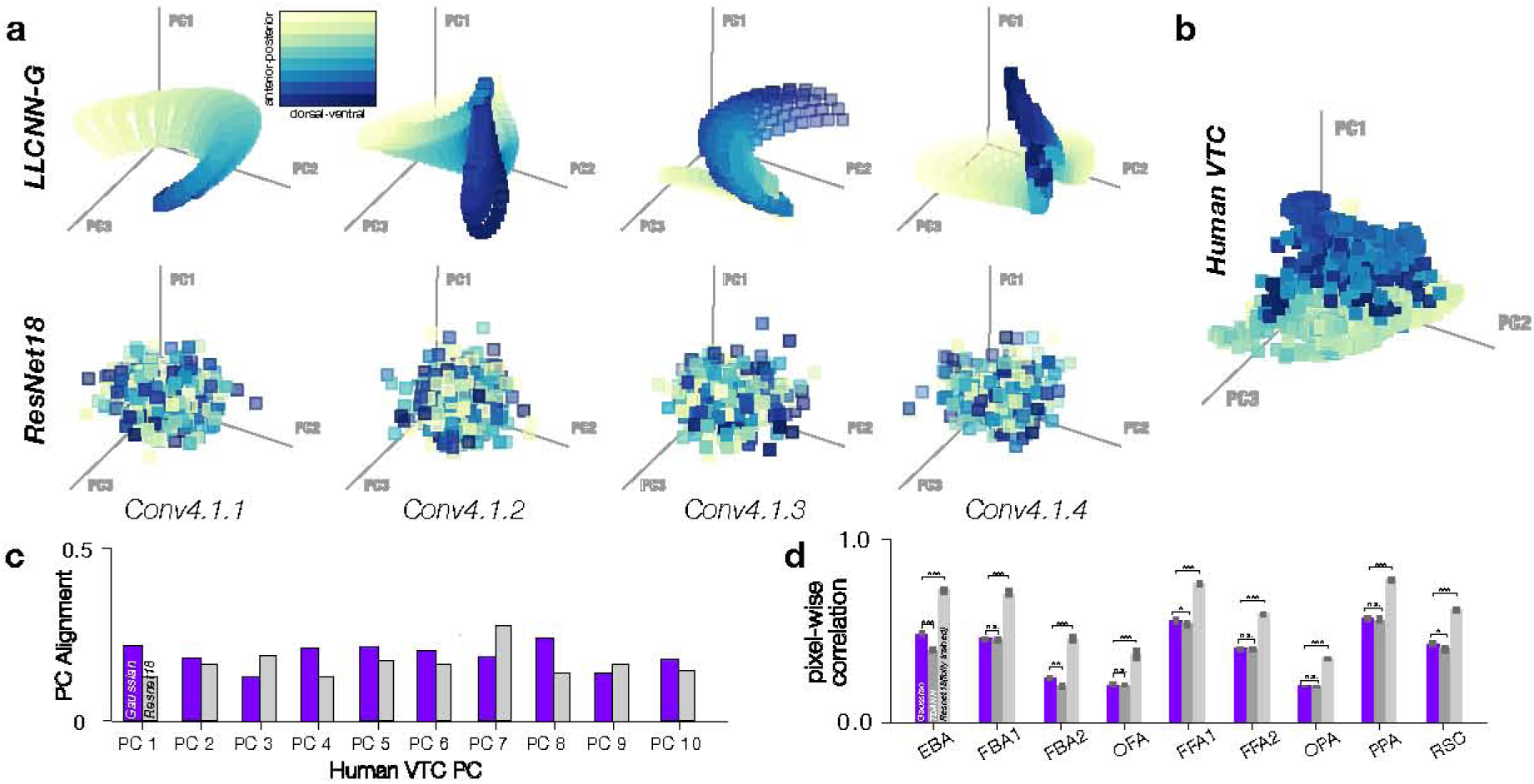
Low dimensional manifold of activity profiles in LLCNN-G model block4 and human ventral temporal cortex. a) Units’ activity profiles are visualized in the top 3 PC space. Each dot corresponds to a unit in the model and units are color coded according to their position on the simulated cortical sheet. LLCNN-G units live on a low-dimensional manifold with maintained physical positions. In contrast ResNet18 units fall on a higher dimensional manifold with no specific position-dependent structure. b) Activity profile manifold similar to (a) but computed over VTC voxels from NSD dataset also exhibit topographical clustering and a low-dimensional manifold. c) Cosine similarity between each pair of model and human brain PCs computed using the NSD dataset. Most top 10 PCs from the human ventral visual cortex are better aligned with the LLCNN-G model compared to ResNet18. d) The unit-voxel correlation between fMRI data from NSD and LLCNN-G model unit activations. Error bars indicate the standard deviation across subjects.

Finally, comparing individual unit activities in each model with voxel activations in the human subjects revealed a higher correspondence (Pearson correlation) between units in the LLCNN-G model and brain in several category-selectivity areas including EBA and FFA1 compared to TDANN [MLF^+^24]. However, correlations were significantly higher for the non-topographical ResNet18 model (Fig. 8d).

## 3 Discussion

Local lateral connections are ubiquitously found across the cortex and a specific pattern of patchy lateral connectivity (aka ”Daisy” patterns) has previously been proposed as a reliable indicator of cortical topography in adult animals of different species [MDCG^+^11]. We incorporated local lateral connections into deep convolutional neural networks and showed that models with lateral connections can closely simulate the topographical organization of neurons across the hierarchy of human visual cortex. The inclusion of lateral connections also yielded an unexpected behavioral outcome: robustness to input perturbations, one of the most remarkable characteristics of sensory processing in humans and other animal species.

### Topographical organization without topography-inducing learning rules and objectives

In the past several years, a number of models were proposed for simulating cortex-like topographical organization in deep neural networks. These models made use of auxiliary learning objectives or learning rules that encourage higher covariance across model units in closer proximity [MLF^+^24, BBP22, LMJ^+^20, FMK^+^23, LDB^+^]. As mentioned in the outset, most recent models rely on strong assumptions about structure and function in biological circuits which challenge their biological plausibility. For instance, approaches that use topography-inducing objective functions [LMJ^+^20, MLF^+^24, BBP22, LDB^+^] implicitly assume that information about synaptic efficacies and axonal lengths are centrally available at the level of each cortical region for the objective function to be rendered computable. On the other hand, approaches that rely on local learning rules such as the self-organizing maps [Koh82], have been shown to lack the capacity to learn rich hierarchical representations similar to those in the human brain [DQBB24]. In contrast, in our work, we show that cortex-like topographical organization can be closely recapitulated by incorporating lateral connections into deep neural networks trained with top-down credit assignment algorithms. This results in a substantially simplified model with more biologically supported assumptions about the neural circuitry that gives rise to the cortical topography. Our topographical LLCNN model extends prior models of cortical topography in early visual cortex [VdM73, Lin86] to the entire ventral visual pathway, suggesting that the same circuit motif (local lateral connections) is key to the emergence of topographical organization in the cortex.

### Top-down Influence on Shaping Cortical Topographical Organization

Most models of cortical topography have traditionally relied on local learning rules or objectives [Koh82, VdM73]. Although many of these models successfully captured various aspects of neuronal organization in the primary visual cortex, their application was largely restricted to early sensory cortices. This limitation likely stems from the inefficiency of local learning rules in constructing the complex representational hierarchies found in modern deep networks trained with backpropagation. Our present work underscores the potential role of top-down credit assignment mechanisms (i.e., backpropagation of error) in shaping cortical topographical organization. By combining local lateral connections with top-down learning via backpropagation, the LLCNN model not only replicated the topographical organization of the ventral visual cortex, but it also did so with significantly less negative impact on the utility of its representations—an essential factor for vision-based tasks such as object recognition. Although the biological plausibility of the backpropagation algorithm remains a subject of ongoing debate, several biologically-aligned variations of this algorithm have been proposed in recent years [AJF^+^22, LSM^+^20], opening new avenues for further exploration.

### Topography as a consequence of a circuit motif that improves robustness

The question of why the cortex is topographically organized has intrigued neuroscientists for decades. The functional significance and evolutionary purpose of cortical columns and their structured arrangements on the cortical surface remain subjects of ongoing debate. While some argue that cortical columns may have no specific computational role [HA05], a widely accepted view suggests that functional modularity arises from a bias toward short connections—a plausible evolutionary strategy [JJ92]. In support of this idea, recent computational models have demonstrated that minimizing cross-unit connections across layers of neural networks leads to the emergence of cortex-like topographical organization [BBP22]. This finding suggests that the orderly arrangement of the cortex may have evolved to optimize limited skull space, thereby minimizing the volume of nerves required to connect neurons across different brain regions.

Our findings offer an alternative, and perhaps complementary, perspective to this theory. We show that learning in locally laterally connected neural networks inherently reduces cross-layer connection volume, even without explicitly penalizing longer inter-neuron connections. More importantly, the formation of neural ensembles within these networks not only leads to spatial clustering but also fosters computational interconnections that significantly enhance robustness to input perturbations. This ensemble-based view of neural computation aligns with well-established machine learning techniques, such as bagging [Bre96], boosting [Fre95], and random forests [Bre01], all of which demonstrate that ensembles of weak processing units can combine to form more powerful and resilient systems for information processing.

In conclusion, our findings highlight the multifaceted role of cortical local lateral connections, showing that they not only optimize inter-neuronal wiring but also play a crucial role in enhancing the robustness of neural representations. Most notably, we demonstrate that the emergence of complex, multi-scale topographical maps can occur naturally, without the need for explicit topography-inducing learning rules. This challenges longstanding theoretical models and assumptions about cortical organization. The discovery that lateral connectivity contributes to both efficient neural wiring and improved computational robustness offers a new functional perspective on cortical topography. This work paves the way for future explorations into the organizational principles of the brain and opens new avenues for designing neural network models that more faithfully replicate the brain’s remarkable efficiency and resilience.

## 4 Methods

### 4.1 Laterally-connected topographic neural networks (LLCNNs)

Architecture. We used the ResNet18 architecture [HZRS16] for all model variants in our work. In contrast to the original ResNet18, we arranged the model units (i.e. convolutional kernels) on a 2D plane that simulated the 2D surface of the cortex (i.e. *simulated cortical sheet* ). The arrangement was systematic both within and across layers of the network, grounding each unit in a physical space that allowed defining physical distance between each pair of units both within a layer and across (Fig. 1), thereby breaking the permutation symmetry not only among units within kernels but also among kernels themselves. This was inspired by the continuous physical proximity of neurons within cortical areas that are hierarchically close (e.g. V1 and V2).

We considered the computations performed by the local lateral connections to be primarily captured by the following equation:

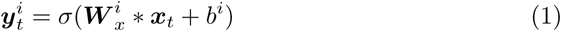

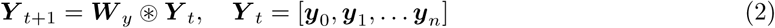

where *x*_t_ and *y^i^* denote the input to a given network layer, and output of the *i*th kernel from that layer respectively. *W* _x_ denotes the kernel weights of the convolution operation ∗ applied along the spatial dimensions of the input, and *W* _y_ the local lateral connection kernel of the Kernel Pooling (KP) operation ⊛ applied along the kernel dimensions of *Y* _t_ [BIDR22, BFCP22]. This approach mirrors the principles of spatial pooling operation but *applied along the channel dimension(s) of the layer activations*. For simplicity, in our simulations, we consider a single-step variation Eq. 1 where the output is computed only once and not iteratively for multiple steps.

To allow embedding of units from different layers within the same simulated cortical sheet, despite the differences in the number of kernels within each layer, we replaced the traditional zero padding with Continuous Padding (CP) which involved appending the size-matched activation from the preceding layer to the current layer’s activation before applying the KP operation. Several variations of the KP models were trained including: 1) Kernel Average Pooling (KAP): Computes the average of unit activations within the region of the feature map covered by the filter (LLCNN-S); 2,3) Kernel Gaussian Pooling, Kernel Mexican-hat Pooling: Computes the weighted average of the unit activation within the filter regions based on a Gaussian and Mexican-hat weighting function respectively (LLCNN-G and LLCNN-MH); 4) Learnable average pooling where the KP parameters *W*_y_ are considered as learnable parameters and are optimized along with other network parameters on minimizing the objective function (LLCNN-L).

### LLCNNs with locally connected layer

We additionally trained a variation of LLC model where we replaced the first convolutional layer of the network with a locally-connected layer. Unlike convolution layers which share weights across spatial positions, locally connected layers employ independent sets of weights at each spatial location, enhancing the expressiveness of the preference map, particularly within V1. However, it’s noteworthy that this modification significantly intensifies GPU memory usage. Therefore, in our implementation, we opt to replace only one layer with a locally-connected layer in each model to manage memory constraints. We used a locally-connected layer with kernel size 3 and stride 1 which divided the model’s full visual field into 56 × 56 patches. Following prior observations reporting the arrangement of neural selectivity according to eccentricity and polar angle [AMSK09], we arranged the layer weights corresponding to different patches on the simulated cortical sheet following a similar pattern (Fig. 2). We converted the Cartesian coordinates of each unit to polar coordinates and subsequently rearranged the units based on their polar positions. In the Cartesian coordinate system, both the x-axis and y-axis range from -168 to 168, positioning the origin as the center of the spatial arrangement.

#### Training

Each neural network model was trained on the Imagenet dataset [DDS^+^09] for 100 epochs. We used the Adam optimizer [KB14] for computing the parameter updates from gradients and a scheduler with an initial learning rate of 0.1. We considered training models with fixed-size (fixed sized-LC) and exponentially decaying (decayed-LC) lateral connection size. In the fixed sized-LC mode, the size of the lateral connection kernel *W* _y_ was predetermined and held constant as 0.1 and 0.23 during training while in the decayed-LC mode, we began training using a maximally-sized lateral connection kernel *W* _y_ which was equal to the size of the layer map and exponentially decayed the size throughout training. Empirically, we observed that the decayed-LC training leads to the emergence of topographical organization at increasingly finer scales.

#### Objective function

We trained the neural network models using supervised or unsupervised learning objectives. The preferred supervised networks were solely trained to minimize the object classification cross-entropy loss on the ImageNet dataset. For the unsupervised training, we considered a contrastive approach, Barlow Twins, which has been shown to produce rich visual representations across various architectures [ZJM^+^21]. We do not believe that different types of supervision make a significant difference in the eventual topography as our results from supervised and unsupervised learning yield similar topographic organization.

### 4.2 V1 analyses

#### Preference map

For analyses presented in Fig. 2, we quantified the orientation, spatial frequency and chromaticity of selectivity of units in layers of each neural network model. We used a stimulus set consisting of grating stimuli of various orientations, spatial frequencies and colors (i.e. black/white vs. red/cyan) [MLF^+^24]. For each unit, we identified the orientation, spatial frequency, and color that elicited the highest response in that unit and considered that as the preferred stimulus property within each category. To visualize the preferred stimulus property maps, we assigned a color to each unit in the map that indicated the preferred stimulus, which maximizes unit activation strength. The unit size encodes the stimulus selectivity. To quantitatively measure orientation selectivity, we employed the circular variance as in Ringach et al. [RSH02].

#### Pairwise preference difference

For the analysis presented in Fig. 2 and 3, we quantified the change in unit preferences as a function of pairwise distance between units following previous work [MLF^+^24]. For this, we first quantified the preferred stimulus value for each unit (4.2) and then calculated all pairwise unit preference differences between all units in each layer. We divide the difference values by the chance value obtained by random resampling of unit pairs. In this fashion, Δ preference equals 1 indicating the value expected by chance. We then plot the average absolute difference in pairwise stimulus preferences as a function of the pairwise distances on the simulated cortical sheet.

#### Pairwise correlation

Prior work quantified the rate of pairwise neural correlations as a function of cortical distance between recording sites which shows an exponentially decaying pattern across different brain areas [LMJ^+^20]. To replicate this in the model, we quantified the pairwise correlation between units within each model layer across all stimuli in the NSD dataset. We then computed the average pairwise Pearson correlation between model units within each layer and plot those values against the pairwise distances on the simulated cortical sheet. In contrast to the pairwise preference difference analysis which solely compares the unit activity at their maximum activation for a specific stimulus type, this analysis comprehensively compares entire patterns of unit activity in response to all types of stimuli.

#### Distribution of orientation differences

It is known that the orientation selectivity of 60-75% of laterally connected neurons falls within 45^◦^ of the selectivity of the source neuron [MDCG^+^11]. To investigate this in our model, we calculated the distribution of delta orientation selectivity between each unit and its neighboring units within each neural network layer. We then plotted a histogram of those values and counted the percentage of units with difference in orientation selectivity *<* 45◦.

#### Orthogonality

As previously demonstrated [NNDC12], orientation and spatial frequency gradient directions on the surface of cortex are highly skewed towards orthogonality. To quantify this in our model, we first mapped the contours of orientation and spatial frequency preference maps within each model layer. We then superimposed the contour maps of the two attributes and identified the cross sections and quantified the angles between them using Python OpenCV package. Finally, we visualized the distribution of angles at the intersections of these maps.

#### Pinwheel density

Each unit was assigned a ’winding number’ [MLF^+^24], which was computed by analyzing the preferred orientations of the eight surrounding pixels. A high winding number denoted a clockwise pinwheel, while a low winding number indicated a counterclockwise pinwheel. Pinwheel density was then calculated as the absolute number of pinwheels per square of the cortical column spacing.

#### V4 analyses Best stimulus set

We determined the inputs from MEI [WSF^+^18], contour and shape datasets which elicited the maximum units activations, and visualized them in the simulated cortical sheets with units’ physical positions.

### 4.3 VTC analyses

#### Selectivity map

We evaluated the model units in terms of their selectivity to faces, body parts, scenes, word forms, no-man’s land, and other objects. For this we used the functional localizer stimulus set (fLoc) [SWGS15] to quantify similarity to all categories except no man’s land which was quantified on the stimulus set from [BSMT20] (shared privately by the authors). In addition to selectivity to these categories, we also quantified the selectivity to animate and inanimate objects as well as object real-world size using [KC13].

For each analysis, selectivity was measured by computing the t-statistic [MLF^+^24], where we measured the difference of responses to the target category in contrast to other categories (e.g. face vs. other objects) and normalized the difference by amount variance in each distribution. In this way, the higher the t-statistic was between two distribution of responses the larger and more significant was the difference between the two distributions. Accordingly, higher t-values could be interpreted as a stronger selectivity of units for a particular class of objects.

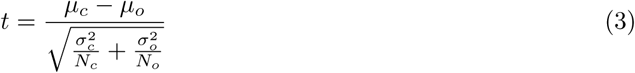

where *µ*, *σ*, *N* denote the average, standard deviation, and the number of samples respectively within the target category *c* and other categories *o*. We then visualized the t-values for all units overlaid on the map of simulated cortical sheet where each unit’s color and size both denoted the selectivity of that unit towards the target category.

#### Smoothness

To compute the smoothness of topography, we define the smoothness score by the difference between the *x*_0_ and minimum correlation values in the pairwise correlation plot normalized by the value at *x*_0_.

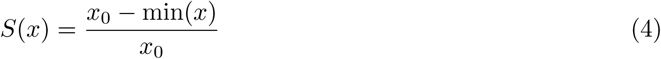

#### Elongation analysis

Most category-selective patches in human cortex are elongated along the posterior-anterior axis of the temporal cortex [NLD^+^11, BSMT20]. We qualitatively observed that when training LLC networks with progressively decaying lateral connectivity (decayed-LC), the corresponding category-selective patches in the model also exhibit elongation along the posterior-anterior axis (shallow to deep layers) of the model. To quantify this phenomenon, we evaluated the elongation of patches at different rates of decay during training. We defined the patch elongation ratio as the length of the patch measured along the posterior-anterior axis divided by the length of patch along the lateral direction which was defined as the axis orthogonal to the posterior-anterior axis.

#### Patch size analysis

To quantify the relationship between the patch size and local lateral connectivity size, we evaluated the size of category patches in models with different kernel pooling (KP) sizes. Using a t-value threshold of 5, we determined the size of candidate patches in the selectivity map by counting the number of activated units within contiguous regions. We then plotted the estimated patch size as a function of KP size.

#### Model-brain patch similarity analysis

We first identified all category-selective patches in the model. To do this, we computed the category-selectivity t-statistic for each model unit in each layer. We classified each unit with a t-value greater than 5 as category-selective and considered any continuous cluster consisting of more than 20 units as a patch. We then computed the cross-validated Pearson correlation between each unit’s average activation across all units within the model patch and the average fMRI response across voxels in the corresponding cortical patch when both were activated by the same stimuli from the Natural Scene Dataset (NSD) [ASYW^+^22]. To evaluate consistency, we calculated the average of all pairwise correlations between the mean activations of corresponding patches across subjects. Specifically, for each patch in a given subject (subject i), we computed its correlation with the same patch in every other subject (subject j), and this process was repeated across all 8 subjects in the NSD. By averaging these pairwise correlations, we obtained a measure of how consistently the patch activations are represented across subjects. This approach provides insight into the reliability and stability of activation patterns across individuals. To assess the statistical significance of the differences of correlations between the two models, a t-test was conducted comparing all pairwise correlations from each model. The resulting p-values indicate the level of significance, with asterisks denoting thresholds: * for p ¡ 0.05, ** for p ¡ 0.01, and *** for p ¡ 0.001. When p-values exceed 0.05, the result is marked as n.s. (not significant), indicating no meaningful difference between the models at that threshold. This test provides a robust measure of whether the observed differences in correlations are statistically significant.

#### Kruskal–Wallis test over 2 sets of t-values

We first calculated two sets of absolute t-values, comparing two categories (i.e., big animacy vs. small animacy, and big inanimacy vs. small inanimacy). Subsequently, we applied the Kruskal–Wallis test to assess statistical differences between the two sets of t-values.

#### Overlapping Area

By comparing the Animacy and Size preference maps, we quantified the number of units located in the overlapping areas between two selective regions (i.e., size-selective and place-selective, as well as size-selective and face-selective regions).

### 4.4 Intermediate Patch Analysis

#### Unit correlation map

To create the unit correlation maps, we calculated the Pearson correlation between each model unit’s activity and the corresponding fMRI responses within the target patch to the full set of stimuli from the NSD dataset. We then visualized the unit correlations using a heatmap plot where the degree of correlation was displayed by the assigned color to each unit.

#### Intermediate patch correspondence

In evaluating topographical models in terms of their alignment with high level visual cortex, these models are commonly inspected for existence of category-selective unit clusters within their layers. While essential, this type of analysis is restricted to known category-selective patches and does not probe whether the topography of these models matches that of the brain outside of these patches. To expand our assessment, we employed an intermediate patch analysis which aims to demonstrate the model’s capability to predict the unknown intermediate patch between two category patches in the visual cortex. To perform this analysis we chose two highly consistent category selective brain regions, namely the Fusiform Face Area (FFA) and Parahippocampal Place Area (PPA) in each subject’s brain. We then found the center voxel of each patch in that subject and mapped the straight line that connects the two points across the surface of the cortex. We then identified the middle point on that line between the boundaries of the two category-selective regions and defined an area of 10 by 10 voxels around that point as the intermediate patch. We followed a similar procedure in the model to identify the model’s intermediate patch and computed the correlation between the two corresponding intermediate patches in the model and the brain in each model.

### 4.5 Behavioral performance, robustness, and other measures

#### Object recognition performance

We evaluated our model’s object recognition performance (top-1 accuracy) using the ImageNet validation dataset [DDS^+^09], which contained 50,000 labeled images distributed across 1,000 classes. The preprocessing routine for these images involved standard transformations: resizing each image to 256 × 256 pixels and then extracting a center crop of 224 × 224 pixels.

#### Quantifying wiring length

To quantify wiring length, we adopted the approach from [MLF^+^24] where units with the highest responses in each layer were first identified and subsequently, the length of inter-layer connectivity necessary to link these identified units were calculated according to their physical distances on the simulated cortical sheet. Specifically, for a given stimulus, we pinpointed the top 5% most responsive units in each of the four adjacent sub-layers. The inter-layer connectivity was established using the k-means clustering algorithm, with connections continuously added until the total ”inertia” of the k-means clustering fell below a specified threshold. The total wiring length was then determined as the sum of the lengths of each individual inter-layer connectivity and the intra-layer connectivity which connected the centroids across layers. It is important to note a distinction from a previous study [MLF^+^24], where the average wiring length across all shift directions was reported. In our model, the anterior and posterior directions were explicitly defined, eliminating the need for direction shifting in our analysis.

#### Robustness

We tested the model’s robustness to input perturbations by evaluating the model’s performance under the AutoAttack [CH20] adversarial attack with L2 perturbation epsilon ranging from 0 to 2.5. AutoAttack is one of the most difficult adversarial attacks which consists of a suite of 4 white-box and black-box attacks. To measure the relationship between robustness and laterally connectivity size, we evaluated the model’s performance with varying lateral size under AutoAttack. Because of the high computational cost of this attack, we chose to evaluate the performance of each model using 500 randomly selected samples from Imagenet validation set.

#### Effective dimensionality

we examine the responses of the entire population of units in the layer 4 to a set of 10,112 natural images from the NSD dataset. We summarize the dimensionality of the responses by the effective dimensionality [MLF^+^24].

#### Function manifold

We applied Principal Component Analysis (PCA) to the transposed activations matrix from layer 4 responding to NSD dataset and projected the activation of each unit onto the PCA space. Subsequently, we projected the manifold onto the first three principal components (PC1, PC2, and PC3) to determine the activation values of each unit in response to the stimulus along these directions.

#### PC alignment

The principal component (PC) alignment matrix was computed by evaluating the cosine similarity between each pair of PCs from the LLCNN-G model and human transposed activations. The alignment for each PC was determined by identifying the maximum cosine similarity between the human PC and the model PC. The human PCs were obtained by performing principal component analysis (PCA) on the fMRI data from all voxels in the IT regions.

#### Pixel-wise correlation

We calculated Pearson correlation coefficients for all pairs between unit activations from the entire population in layer 4 and voxel activations from specific selective regions (e.g., EBA) in response to the NSD dataset. The cross-validated maximum correlation from these pairs was then reported as the pixel-wise correlation.

## Acknowledgement

We thank Drs. Nancy Kanwisher, Daniel Yamins, Christopher Pack, Stuart Trenholm, and Arjun Krishnaswamy for their helpful discussions and feedback on the manuscript. We also thank Drs. Pinglei Bao and Doris Tsao for sharing the stimulus set from *Bao et al. 2020, Nature* and Dr. Andreas Tolias for sharing the stimulus set from *Willeke et al. 2023, BioRxiv*. This research was supported by the Healthy-Brains-Healthy-Lives startup supplement grant, the NSERC Discovery grant RGPIN-2021-03035, and CIHR Project Grant PJT-191957. P.B. was supported by FRQ-S Research Scholars Junior 1 grant 310924, and the William Dawson Scholar award. All analyses were executed using resources provided by the Digital Research Alliance of Canada (Compute Canada) and funding from Canada Foundation for Innovation project number 42730.

---

**Figure S1:**
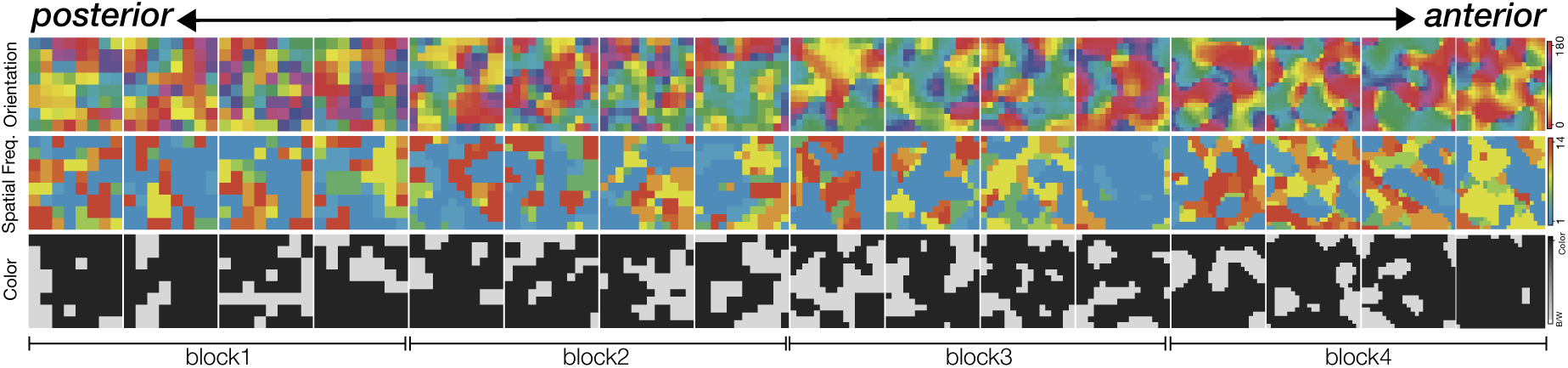
V1 Topography in LLCNN-S model. Selectivity maps to orientation, spatial frequency and color smoothly change across the spatial dimensions of the simulated cortical sheet, both within and across layers. Selectivity maps is shown for all units in all layers of the model. Early to late layers of the model are shown from left to right (posterior to anterior).

**Figure S2:**
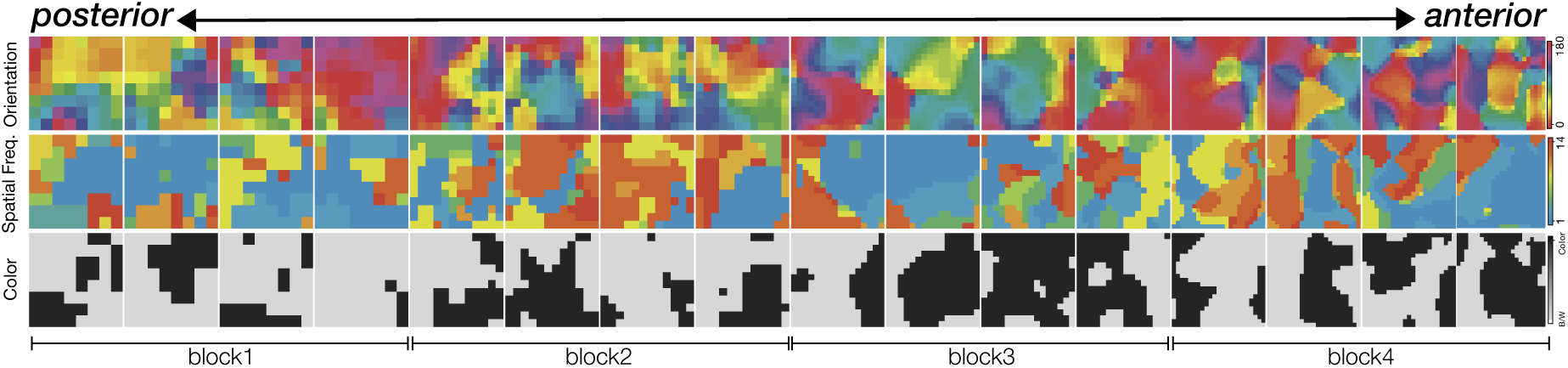
V1 Topography in LLCNN-MH. Selectivity maps to orientation, spatial frequency and color smoothly change across the spatial dimensions of the simulated cortical sheet, both within and across layers. Selectivity maps is shown for all units in all layers of the model. Early to late layers of the model are shown from left to right (posterior to anterior).

**Figure S3:**
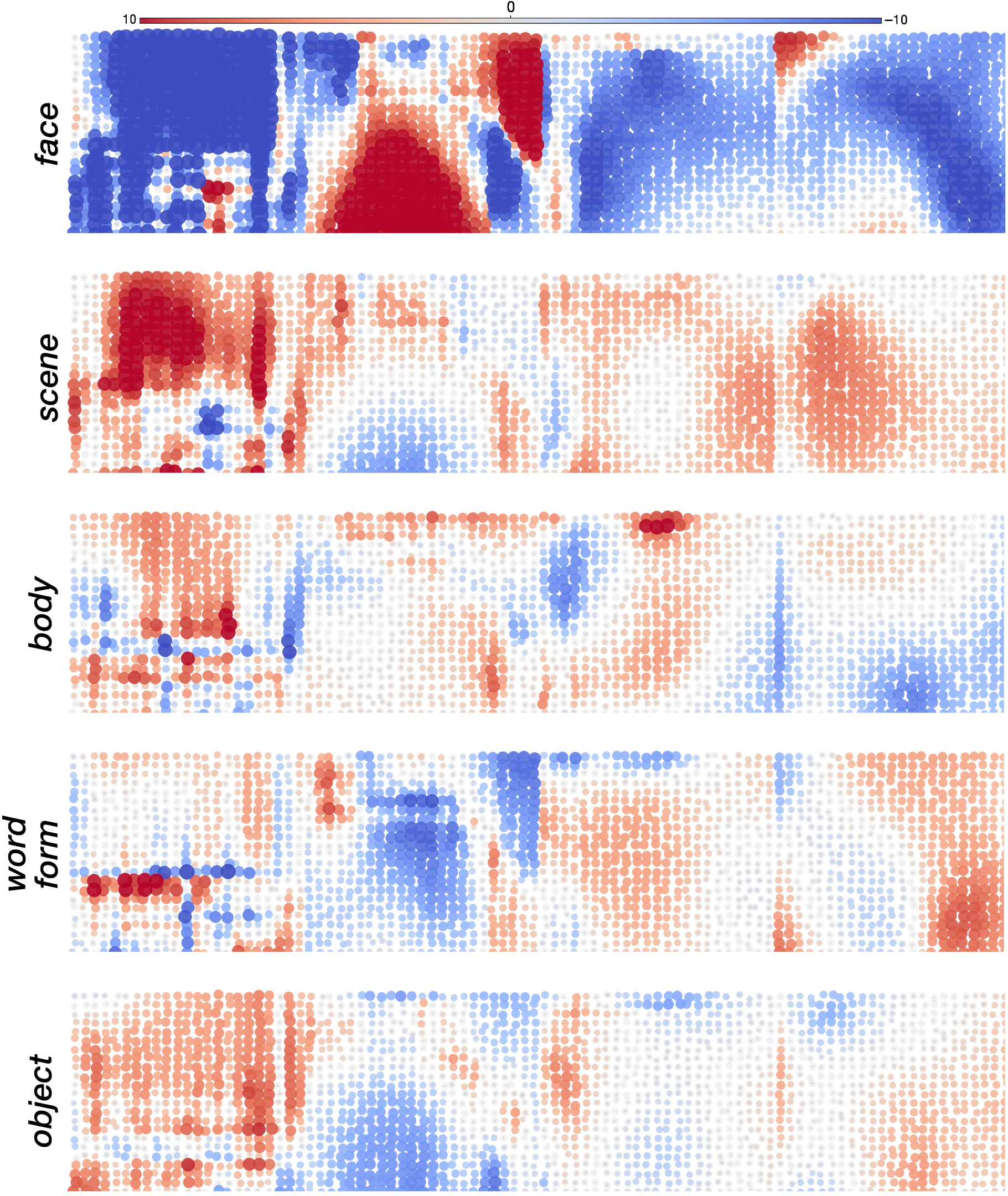
Topographical arrangement according to object categories in deep layers of LLCNN-S model. Unit responses were assessed concerning their selectivity to five distinct categories of images, namely face, scene, body, characters, and objects. Selectivities are shown for the 4 convolutional layers in block 4 of the ResNet18 architecture. Color denotes the selectivity of each unit to each category.

**Figure S4:**
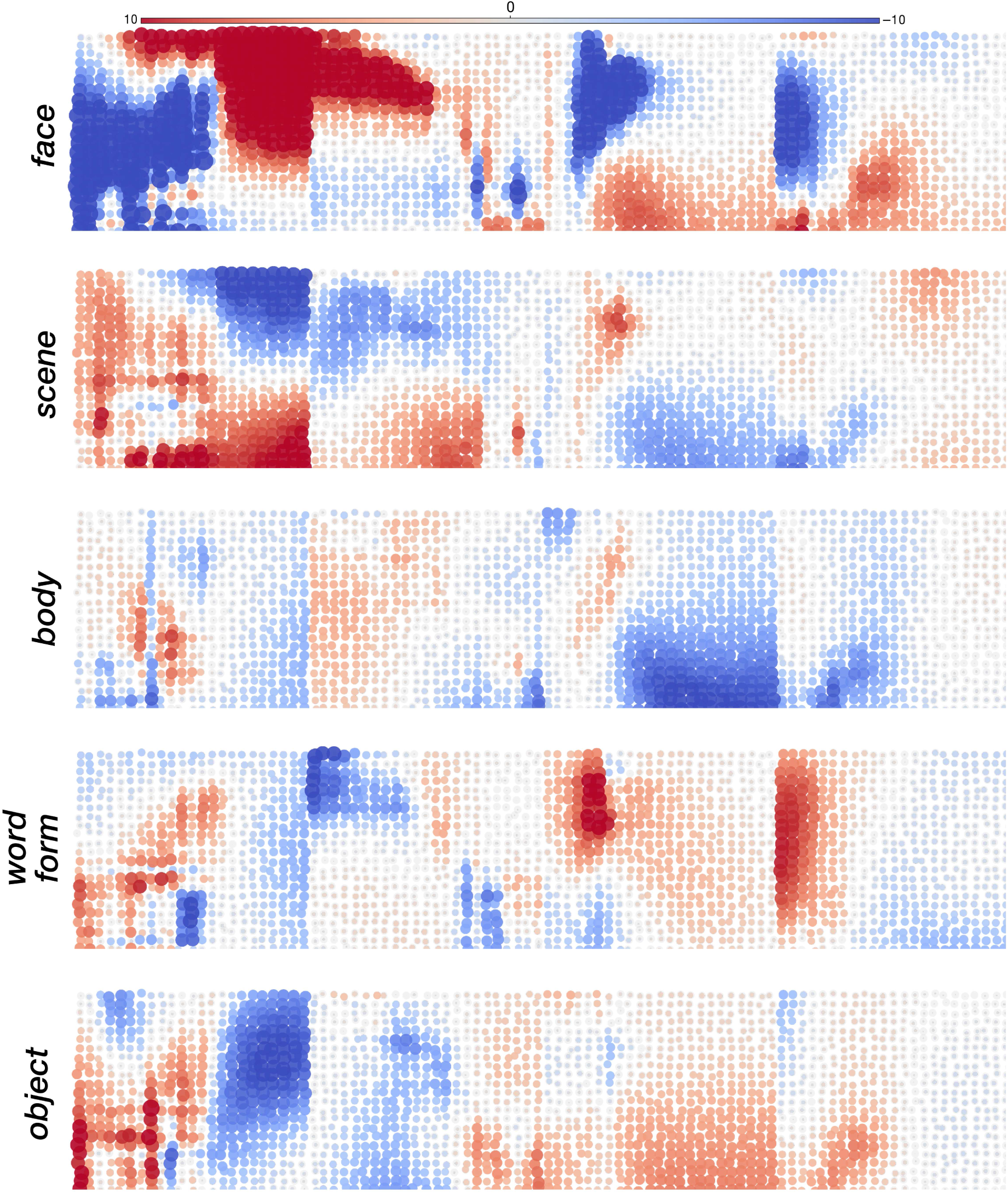
Topographical arrangement according to object categories in deep layers of LLCNN-MH model. Unit responses were assessed concerning their selectivity to five distinct categories of images, namely face, scene, body, characters, and objects. Selectivities are shown for the 4 convolutional layers in block 4 of the ResNet18 architecture. Color denotes the selectivity of each unit to each category.

**Figure S5:**
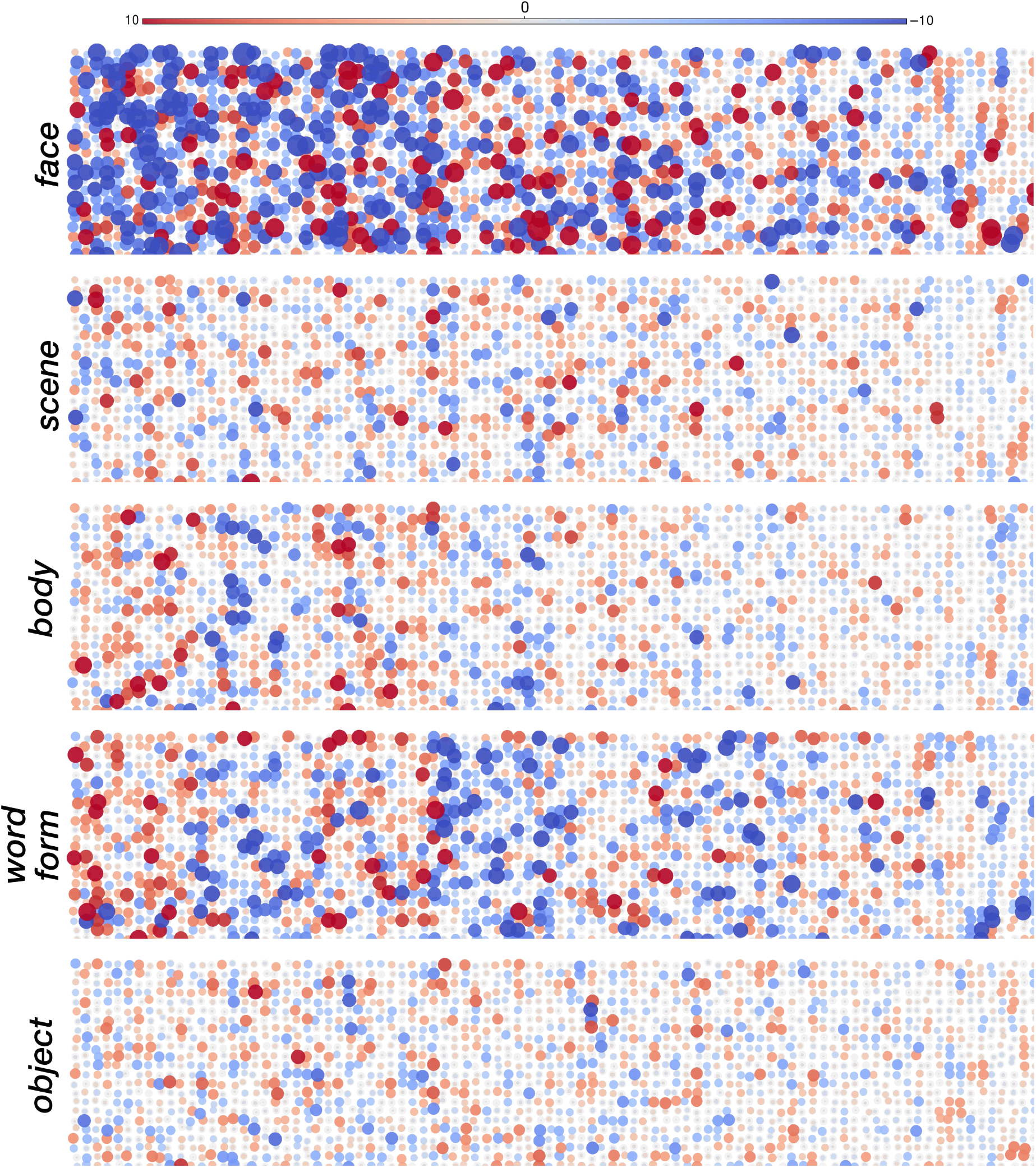
Topograhical arrangement according to object categories in deep layers of LLCNN-L model. Unit responses were assessed concerning their selectivity to five distinct categories of images, namely face, scene, body, characters, and objects. Selectivities are shown for the 4 convolutional layers in block 4 of the ResNet18 architecture. Color denotes the selectivity of each unit to each category.

**Figure S6:**
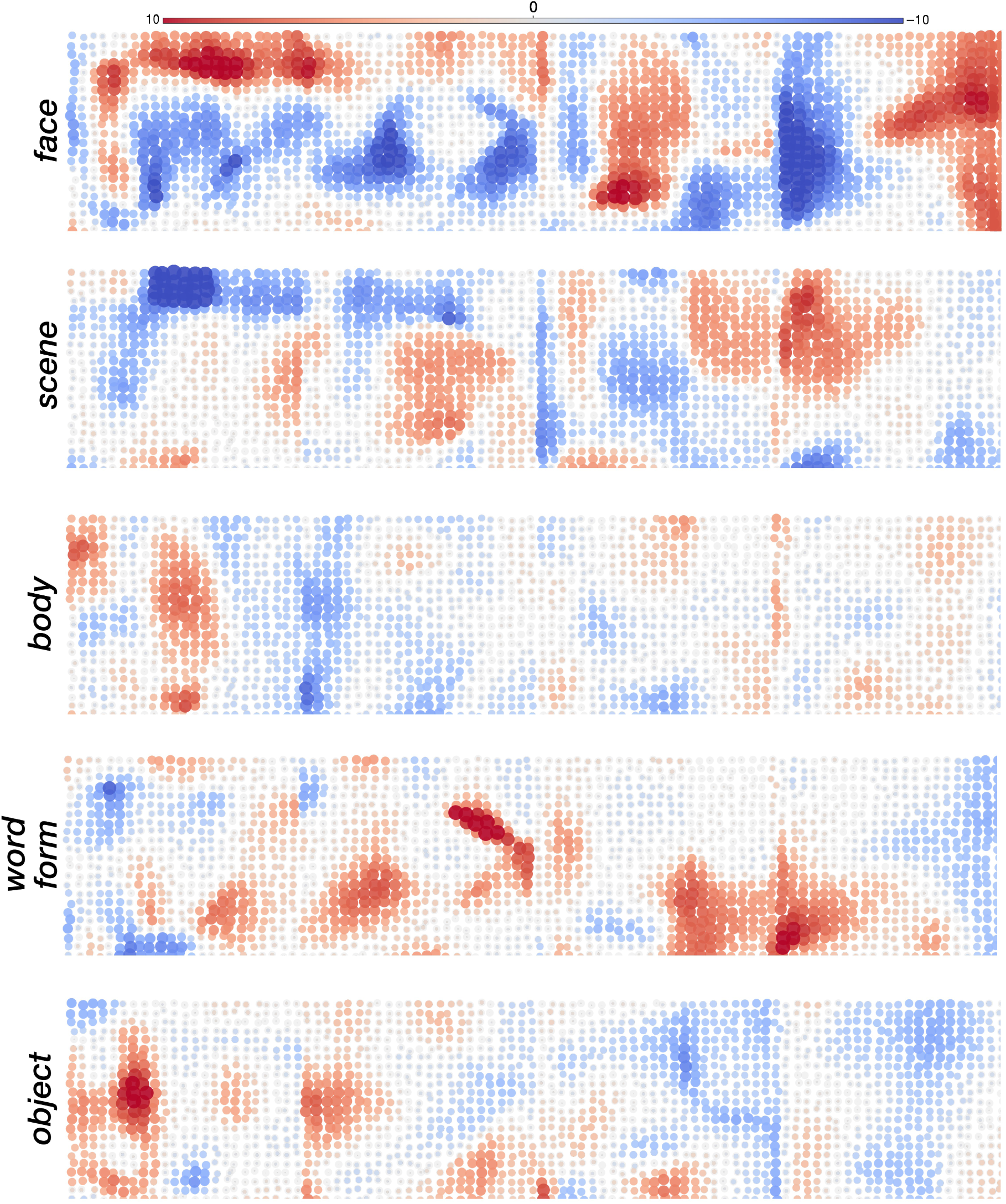
VTC-like topography in LLCNN model trained unsupervised via Barlow’s Twins [ZJM^+^21]. Unit responses were assessed concerning their selectivity to five distinct categories of images, namely face, scene, body, characters, and objects. Selectivities are shown for the 4 convolutional layers in block 4 of the ResNet18 architecture. Color denotes the selectivity of each unit to each category.

**Figure S7:**
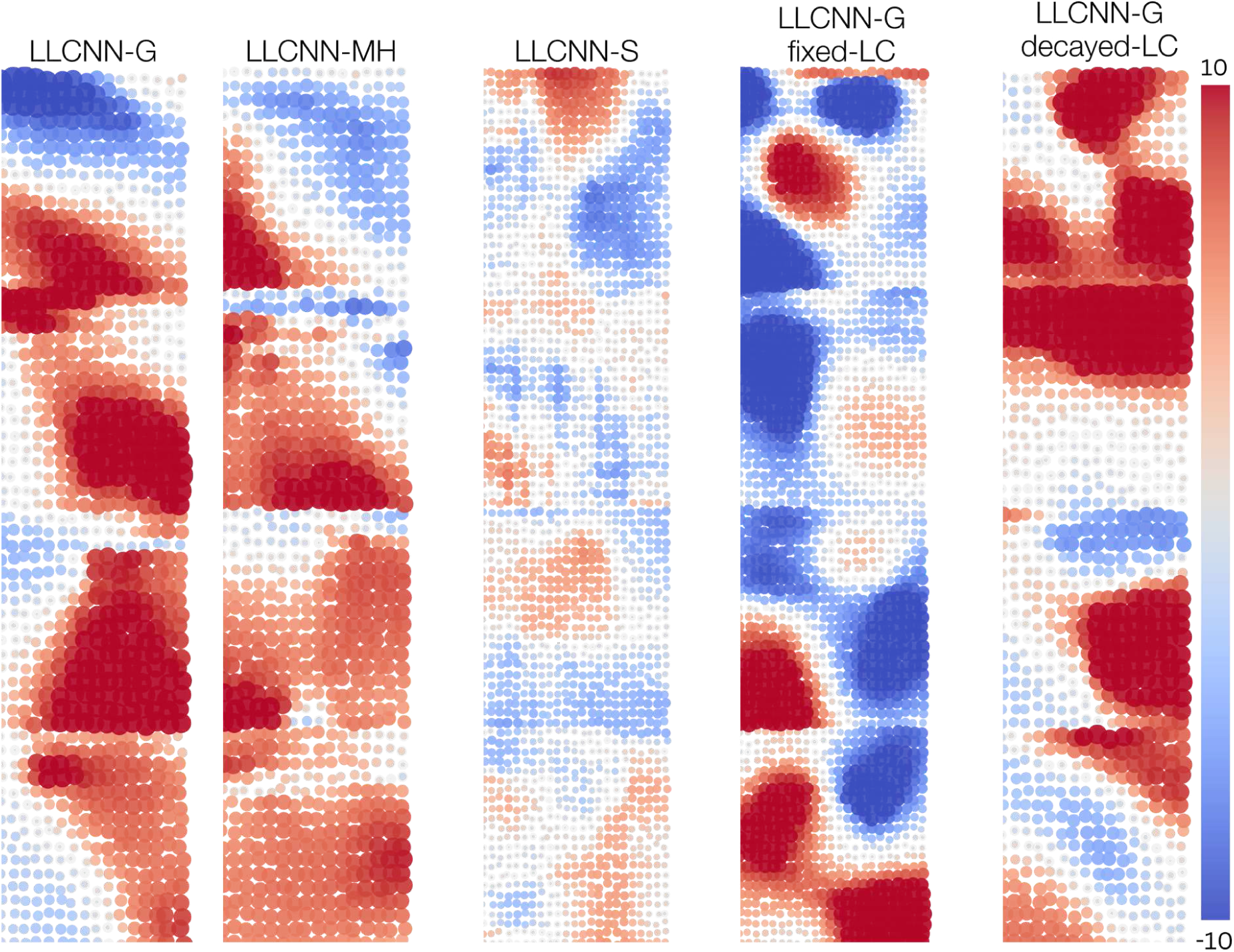
Face selectivity in LLCNN model across different model variations and training regimes. Continuous face-selective patches emerged in all variations of the LLCNN model. Face selectivity index was visibly lower in the LLCNN-S model, possibly because of its lower performance on Imagenet categorization task. Training with fixed lateral connections (fixed-LC) led to more blob-like category selective patches and more discontinuities in category-selectivity across layers compared to the version with exponentially decaying lateral connectivity size (decayed-LC).

**Figure S8:**
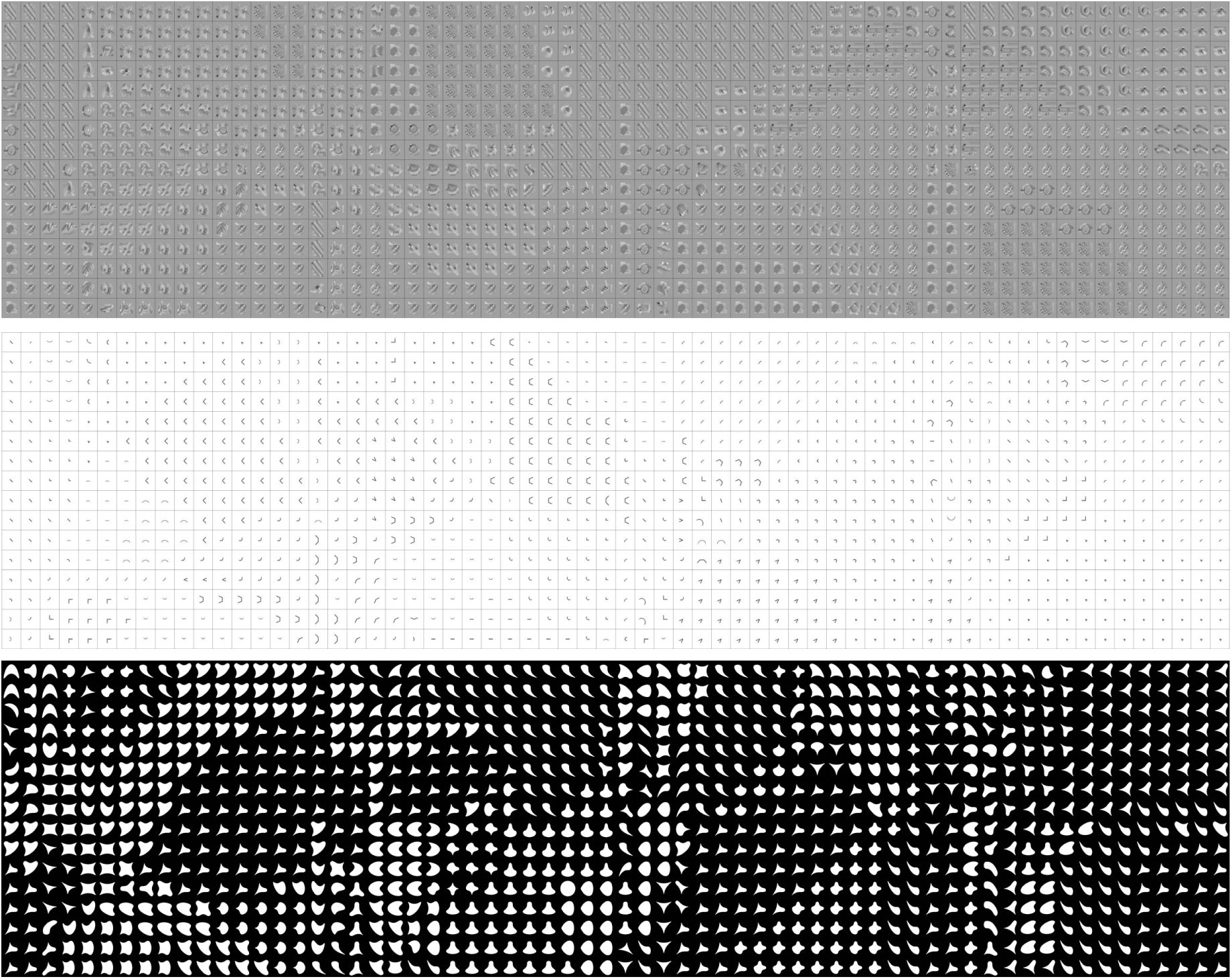
A map of best stimuli for units in 4 consecutive layers of block3 in LLCNN-G model. The three rows correspond to stimuli from [WRF^+^23], [JALT21], and [PC02] respectively. Model units are clustered into groups for each stimulus set.

**Figure S9:**
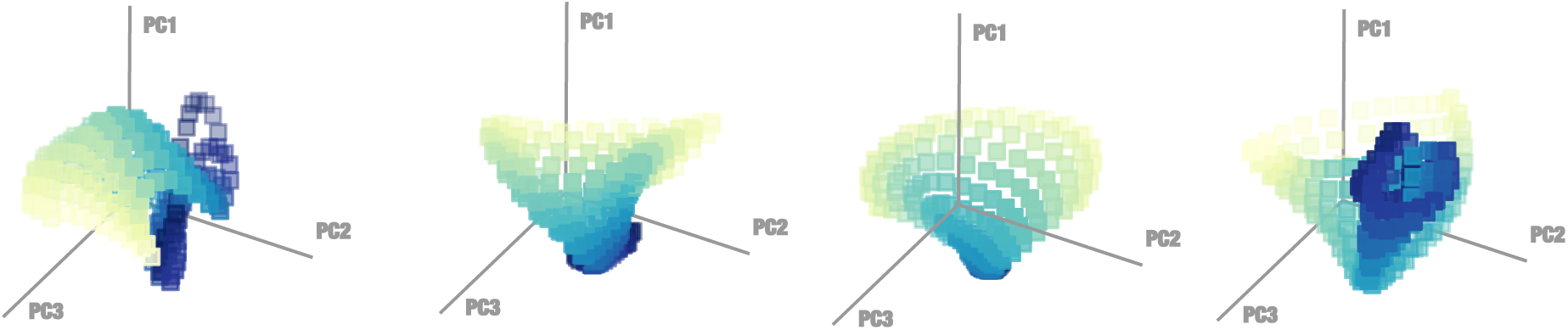
Low dimensional manifold of activity profiles for block 3 of LLCNN-G model. Plots are similar to those in Fig. 8a but for block 4 of the LLCNN-G model. The activity profiles lie on a low dimensional manifold that preserves the spatial arrangement of the units on the simulated cortical sheet as well.

## Footnotes

1 We compared the visual categorization performance with the self-supervised version of TDANN as only this model variation (and not their supervised-trained model) exhibited cortex-like topographical organization.

